# Interpretable network-guided epistasis detection

**DOI:** 10.1101/2020.09.24.310136

**Authors:** Diane Duroux, Héctor Climente-González, Chloé-Agathe Azencott, Kristel Van Steen

## Abstract

Detecting epistatic interactions at the gene level is essential to understanding the biological mechanisms of complex diseases. Unfortunately, genome-wide interaction association studies (GWAIS) involve many statistical challenges that make such detection hard. We propose a multi-step protocol for epistasis detection along the edges of a gene-gene co-function network. Such an approach reduces the number of tests performed and provides interpretable interactions, while keeping type I error controlled. Yet, mapping gene-interactions into testable SNP-interaction hypotheses, as well as computing gene pair association scores from SNP pair ones, is not trivial. Here we compare three SNP-gene mappings (positional overlap, eQTL and proximity in 3D structure) and use the adaptive truncated product method to compute gene pair scores. This method is non-parametric, does not require a known null distribution, and is fast to compute. We apply multiple variants of this protocol to a GWAS inflammatory bowel disease (IBD) dataset. Different configurations produced different results, highlighting that various mechanisms are implicated in IBD, while at the same time, results overlapped with known disease biology. Importantly, the proposed pipeline also differs from a conventional approach were no network is used, showing the potential for additional discoveries when prior biological knowledge is incorporated into epistasis detection.

## 1 Background

Genome-wide association studies (GWAS) have identified over 70 000 genetic variants associated with complex traits [5]. Often these variants altogether do not explain the whole variance of a trait. A representative example is inflammatory bowel disease (IBD), like Crohn’s disease and ulcerative colitis. Pooled twin studies estimate their heritabilities at 0.75 and 0.67 respectively [15]. Yet, despite large GWAS that identified over 200 IBD-associated loci [11], a low proportion of their variance has been explained [38]. Possible explanations include a large number of common variants with small effects, rare variants with large effects not covered in GWAS, unaccounted gene-environment interactions, and genetic interactions [29]. In this article we explore the latter, called epistasis, which has been linked to IBD in the past [14, 26, 30, 33, 45, 53]. Often, two types of epistasis are described: biological and statistical epistasis [31]. Broadly described, biological epistasis refers to a physical interaction between two biomolecules that has an impact on the phenotype. Statistical epistasis refers to departures from population-level linear models describing relationships between predictive factors such as alleles at different genetic loci.

Genome-wide association interaction studies (GWAIS) focus on the detection of statistical epistasis. To date, these studies have produced few replicable, functional conclusions, and specific gene-gene interactions have rarely been identified. This may be due to the small effects sizes of the interactions, the low statistical power, or the absence of a widely accepted GWAIS protocol. Even in the absence of statistical challenges, GWAIS are usually conducted on single nucleotide polymorphisms (SNPs), and SNP-interactions often lack a straightforward functional interpretation. Moving from SNP- to gene-level tests, which jointly consider all the SNPs mapped to the same gene, might address both shortcomings. First, aggregating SNP pair statistics into gene pair statistics is likely to increase the statistical power when dealing with complex diseases [49]. Second, converting statistical findings into biological hypotheses [23] may facilitate their functional interpretability [22].

To both reduce the number of tests and improve the interpretablity of significant SNP interactions, some authors propose examining only pairs of SNPs likely to be functionally related [32]. Such approaches use prior biological knowledge, for instance, of SNPs involved in genes that establish a protein-protein interaction [17]. Yet, limiting studies to one particular kind of gene-gene interaction might be reductive. To tackle that issue, Pendergrass et al. [34] developed Biofilter, a gene-gene co-function network, which aggregates multiple databases. Additionally, such approaches often require as well a proper mapping of SNP to genes.

In this article, we propose guiding the search for statistical epistasis using plausible biological epistasis. Taking exclusively interactions reported from at least 2 different sources in Biofilter, we compile a subset of gene-gene interactions that are biologically plausible. Then, we exclusively search for those interactions in a GWAIS dataset, reducing the multiple test burden and improving the interpretability. We investigate different ways of mapping SNPs to genes and use the adaptive truncated product method [39] to estimate the association of gene pairs. Network and pathway analyses are used to further assist in the interpretation of epistasis findings. The proposed pipeline is applied to GWAS data from the International IBD Genetic Consortium [11].

## 2 Data description

We investigated the IIBDGC dataset, produced by the International Inflammatory Bowel Disease Genetics Consortium (IIBDGC). This dataset was genotyped on the Immunochip SNP array [8]. We performed quality control as in Ellinghaus et al. [12], hereby reducing the number of SNPs from 196 524 to 130 071. The final dataset contains 66 280 samples, out of which 32 622 are cases (individuals with IBD) and 33 658 are controls. The large sample size of this dataset helps overcoming the issue of reduced statistical power that is common in GWAIS.

The IIBDGC dataset aggregates different cohorts, and contains potentially confounding population structure. As in Ellinghaus et al. [12], we used the first 7 principal components to model population stratification. Because several epistasis detection methods, such as those implemented in PLINK [37], cannot include covariates in their logistic regression models, we instead adjusted the phenotypes by regressing out those principal components. In other words, we derived adjusted phenotypes from the logistic regression model by subtracting model-fitted values from observed phenotype values, i.e. response residuals (see Supplementary Fig 1).

## 3 Analysis

### 3.1 SNP to gene mapping: Chromatin contacts map more SNPs per gene than other mappings

In this article, we present a pipeline to detect gene epistasis across the edges of a network. We extract interacting pairs of genes from the gene-gene Biofilter network to obtain candidate gene epistatic pairs (*gene models*). We considered three ways to match genes to SNPs and obtain *SNP models* from them: *Positional*, *eQTL* and *Chromatin* (detailed in section *From gene models to SNP models*). *Chromatin* produced the largest number of unique SNP-gene mappings (2 394 590), an order of magnitude more than *eQTL* (411 120) and *Positional* (174 879) (Table 4). The *Chromatin* mapping had on average the largest number of SNPs mapped on to a gene, followed by *eQTL* and *Positional* (Fig 1A). Nonetheless, the number of SNPs mapped to a gene varied considerably across genes (Fig 1B). In addition, the number of SNPs mapped to a same gene varied considerably across analyses (Fig 1C, D and E): in general, the genes with most SNPs mapped using the *eQTL* mapping had relatively few SNPs mapped in the *Chromatin* mapping, and vice versa.

**Fig. 1:**
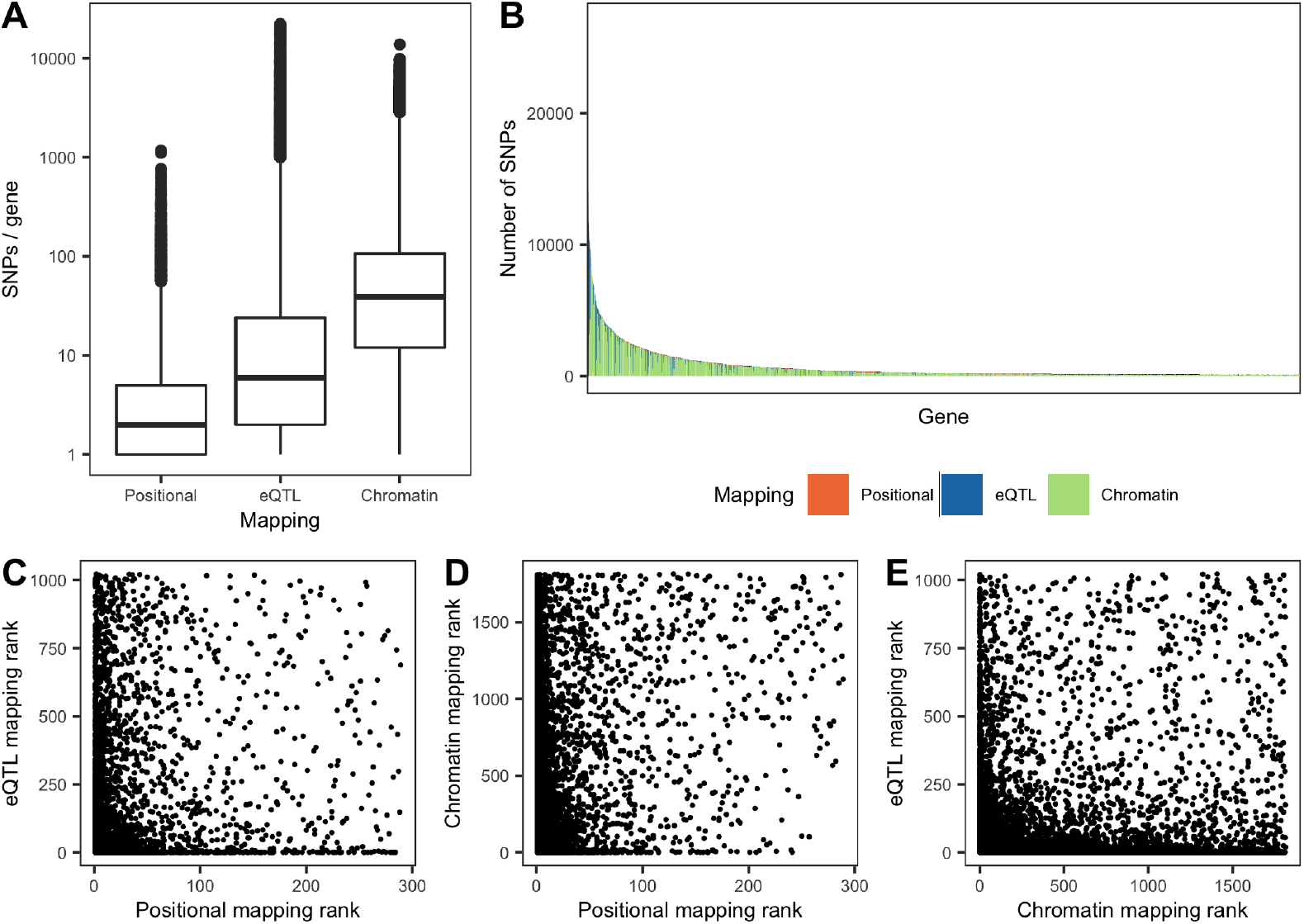
**(A)** Number of SNPs per gene for each of the three mappings described in *From gene models to SNP models*. **(B)** Ranking of genes with most SNPs mapped using any of the mappings, colored by mapping. Only genes with more than 100 SNPs mapped to it are displayed. **(C,D,E)** Comparison between the rank of each gene according to the number of SNPs mapped to it using each mapping.

### 3.2 The *Positional* analysis does not recover any SNP interaction

The aforementioned SNP-gene mappings, and combinations of them (cross-mappings), yielded seven sets of SNP models. Running our pipeline on them resulted in seven epistatic SNP-SNP networks described in Table 1 (for visualization, see Supplementary Fig 2). We also conducted what we called a *Standard* analysis, which reflects a conventional epistasis detection procedure. In this one, we exhaustively searched for epistatic interactions between all the SNPs that passed quality control. Then, we used positional mapping to assign gene interactions to the significant SNP interactions. Strikingly, while the *Standard* analysis generated the largest SNP-interaction network (55 nodes/SNPs and 57 edges/interactions), the *Positional + eQTL* one was the largest by number of interactions (76). The *Positional* analysis produced no significant interactions at all.

**Table 1:** Properties of the SNP networks obtained from different datasets. Nodes are SNPs, which are linked when the SNP model is significant.

| Analysis | SNPs | Edges | Components | Avg. degree |
| --- | --- | --- | --- | --- |
| *Standard* | 55 | 57 | 12 | 2.07 |
| *Positional* | 0 | 0 | - | - |
| *eQTL* | 46 | 64 | 6 | 2.78 |
| *Chromatin* | 20 | 19 | 5 | 1.9 |
| *eQTL + Chromatin* | 44 | 48 | 8 | 2.2 |
| *Positional + eQTL* | 43 | 76 | 6 | 2.7 |
| *Positional + Chromatin* | 21 | 39 | 5 | 1.9 |
| *Positional + eQTL + Chromatin* | 39 | 45 | 6 | 2.3 |

### 3.3 Gene epistasis: “functional” mappings boost discovery and interpretability

Findings of a GWAIS are often presented as a network, with nodes indicating SNPs and edges between nodes being present when the analysis protocol identifies the corresponding SNP pair as significantly interacting with the trait of interest. We converted SNP model networks into gene model epistasis networks (Fig 2), adding an edge between two genes whenever the corresponding gene model significance was ascertained through an ATPM approach. The largest network was obtained under the *Standard* analysis (26 edges, Table 2). The *Positional + eQTL + Chromatin* combinations performed second best (13 edges). Since no significant SNP pairs were detected under *Positional*, no significant gene pairs were produced either. Similarly, the two analyses that included *Positional* information on top of another source barely added any new information in comparison to *Positional + eQTL + Chromatin*: *Positional + eQTL* included a new gene pair (*SPAM1* -*HYAL1*, already detected in the *eQTL* analysis), while *Positional + Chromatin* did not produce any additional pair. On the same line, *eQTL + Chromatin* included only one gene model absent from *Positional + eQTL + Chromatin*: *PLA2G2E-PLA2G2C*. Hence, we removed those three from further analyses, since *Positional + eQTL + Chromatin* captured more biological signal than those separately.

**Fig. 2:**
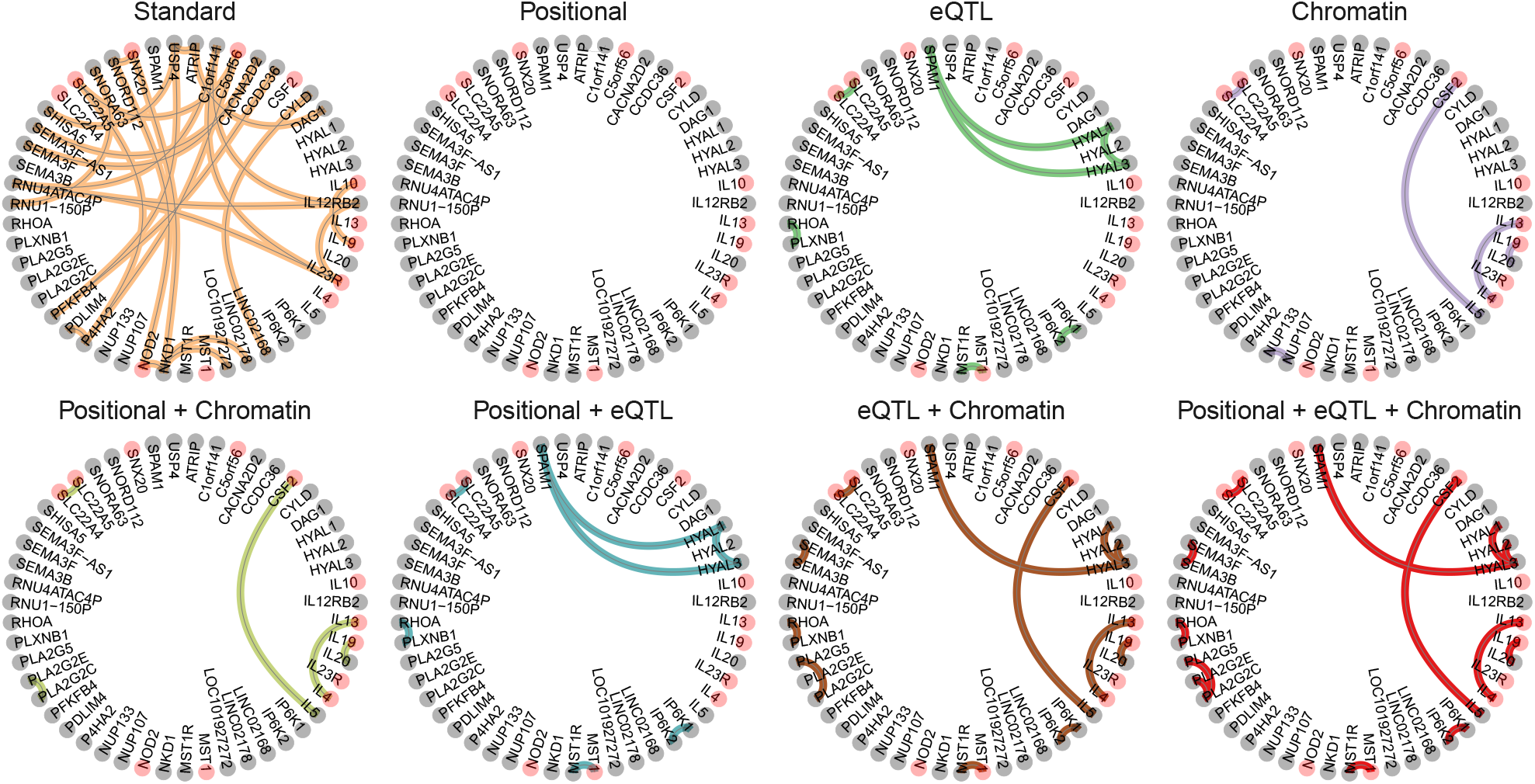
Epistasis networks built from derived significant gene models for the different analysis strategies. Genes associated to IBD in DisGeNET [36] are colored in pink. An alternative layout of the networks is available in Supplementary Fig 3.

**Table 2:** Properties of the gene networks obtained from different datasets. Nodes are genes, which are linked when the corresponding gene model is significant.

| Mapping | Genes | Edges | Components | Avg. degree |
| --- | --- | --- | --- | --- |
| *Standard* | 29 | 26 | 8 | 1.79 |
| *Positional* | 0 | 0 | - | - |
| *eQTL* | 11 | 7 | 5 | 1.27 |
| *Chromatin* | 10 | 5 | 5 | 1 |
| *eQTL + Chromatin* | 22 | 12 | 10 | 1.1 |
| *Positional + eQTL* | 11 | 7 | 5 | 1.3 |
| *Positional + Chromatin* | 10 | 5 | 5 | 1 |
| *Positional + eQTL + Chromatin* | 23 | 13 | 10 | 1.1 |

For both *eQTL* and *Standard* most of the significant SNP models mapped to exclusively one gene model, removing possible sources of ambivalence (Fig 3A). That was less the case under the *Chromatin* analysis, where it was more common for the same SNP model to map to different gene models. We also investigated the relationship between significant gene models and the number of significant SNP models that mapped to them (Fig 3B). Most significant gene interactions were supported by relatively small numbers of SNPs: either few in number, or few with respect to the total number of SNP models for that significant gene model.

**Fig. 3:**
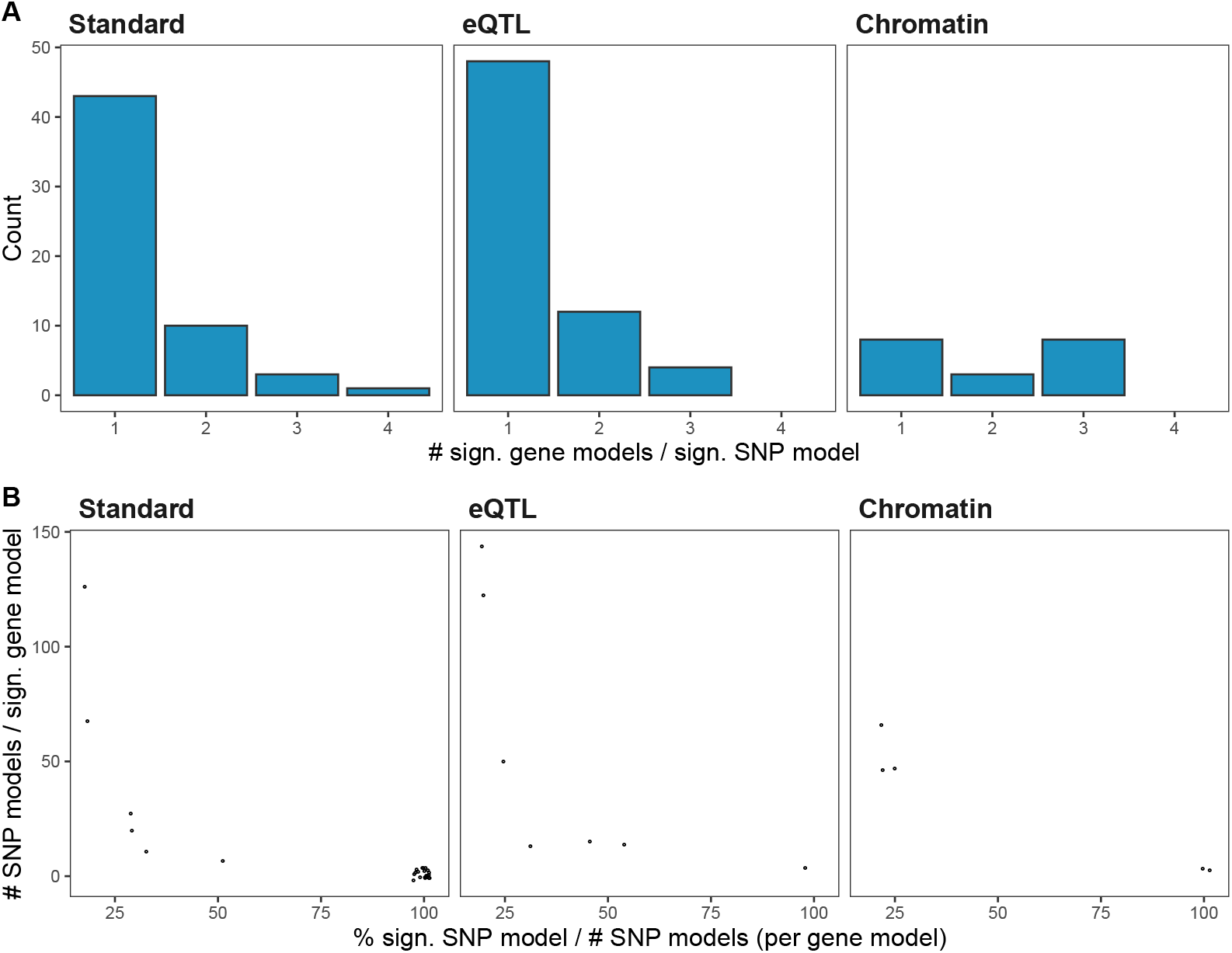
Relationship between the number of significant SNP models and of significant gene models. **(A)** Histogram of the number of significant gene models mapped to the same significant SNP model. **(B)** Relationship between the total number of SNP models mapped to the same significant gene model (y-axis), and the percentage of all the SNP models mapped to the same significant gene model that are significant themselves (x-axis). As multiple points can stack, we introduced a little Gaussian noise on each of them to improve visualization.

### 3.4 Significant SNP pairs are near each other and near loci with main effects

Notably, the SNPs in significant SNP interactions are located near each other in the genome (the median distance between the pair of SNPs in *Chromatin*, *eQTL* and *Positional + eQTL + Chromatin* was 161 kbp). Moreover, they tend to overlap with GWAS main effects loci (Fig 4A). To investigate whether main effects could be driving some of the signals, even when in imperfect LD with epistatic SNP pairs (a phenomenon sometimes referred to as “phantom epistasis” [10]), we conducted a linear regression-based test, including a vector of polygenic risk scores as covariate. The observed effect of many significant SNP model notably decreased when we conditioned on single SNPs in this way (Fig 4B), but not for all. The latter suggests a masking effect opposite to phantom epistasis. However, it is unclear how to adequately correct for multiple hypotheses testing after this adjustment in our setting, and in what follows we still use the unadjusted P-values, with the understanding that some of them may be inflated by weak correlations with main effects.

**Fig. 4:**
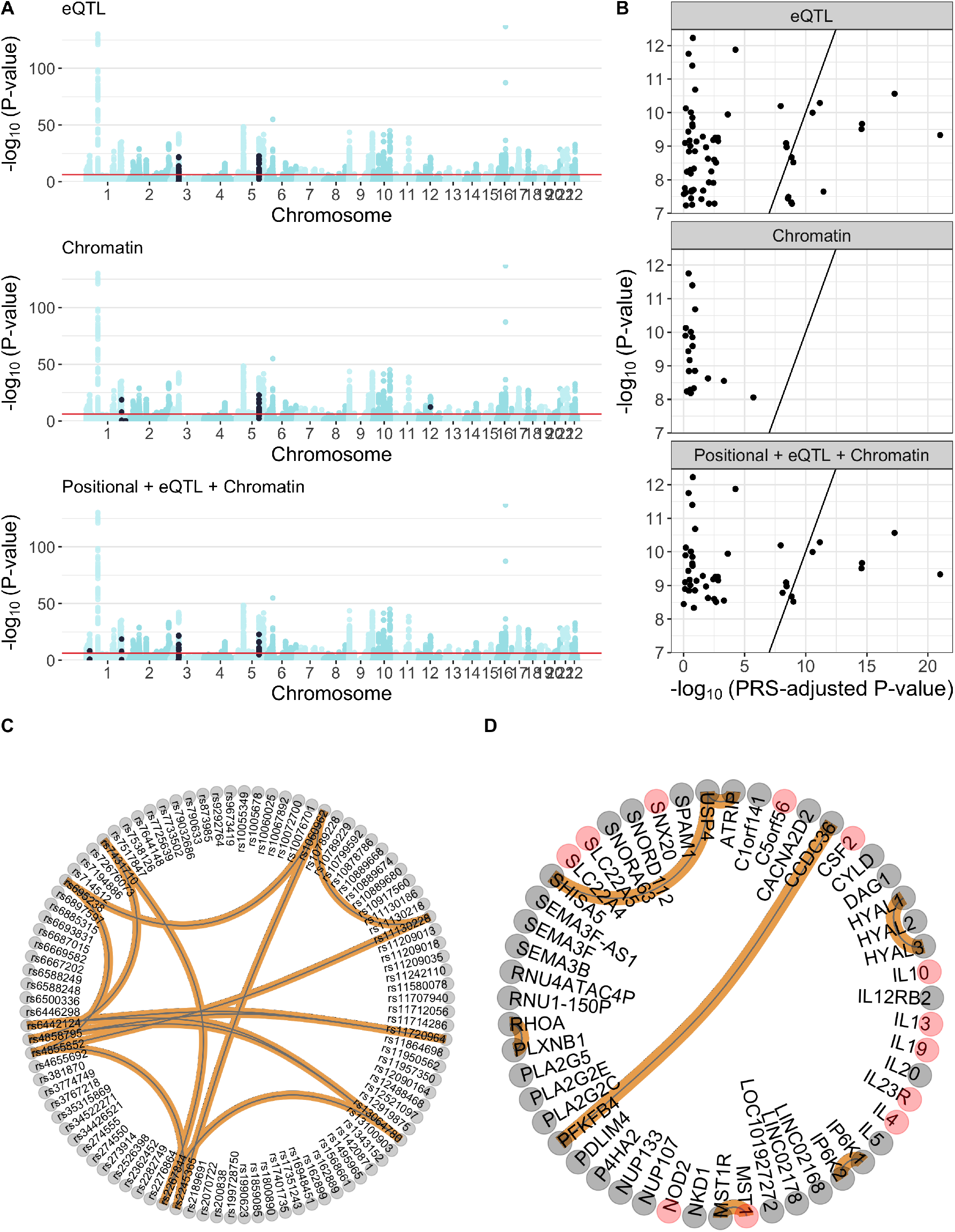
**(A)** Manhattan plot of the main effects, computed using logistic regression. In each subpanel, the SNPs selected via a significant SNP model, by each analysis, are colored in black. For reference, the Bonferroni threshold of main effects significance is displayed with a red horizontal line. **(B)** Comparison between the P-values of the significant SNP interactions, adjusted and unadjusted by main effects (x- and y-axis, respectively). P-values were not adjusted for multiple testing. To help interpretation, we added a *y = x* line. **(C)** Network containing all the SNP models significant in any of the analyses whose P-values after adjusting for PRS were lower than the original P-values. **(D)** Network containing all the gene models significant in any of the analyses that were mapped to one of the significant SNP models from panel C in its corresponding analysis. Genes associated to IBD in DisGeNET [36] are colored in pink.

### 3.5 The type I error is controlled

To evaluate the statistical relevance of the detected gene interactions, we studied if the proposed protocol controlled the type I error. For that purpose, we performed a permutation analysis based on 1 000 permutations for each of the datasets, permuting the phenotypes and running the entire protocol to detect significant gene interactions. This permutation procedure is independent of the one used in the proposed protocol to compute significance thresholds. When at least one significant gene interactions was observed in a permutation, that permutation was considered a false positive (FP). This allowed us to compute the type I error rate as 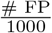. Type I error was under control in all tested experimental settings, with estimates ≤ 6.6% (Table 3).

**Table 3:** Type I error of the protocol presented in *Gene interaction detection procedure*, estimated over 1 000 random permutations, as explained in section *The type I error is controlled*.

| Analysis | Average num. of significant interactions under $H_0$ | Type 1 error (%) |
| --- | --- | --- |
| *Standard* | 0.05 | 3.6 |
| *Positional* | 0.04 | 3.7 |
| *eQTL* | 0.09 | 4.2 |
| *Chromatin* | 0.07 | 6.6 |
| *Positional + eQTL + Chromatin* | 0.04 | 3.6 |

**Table 4:**
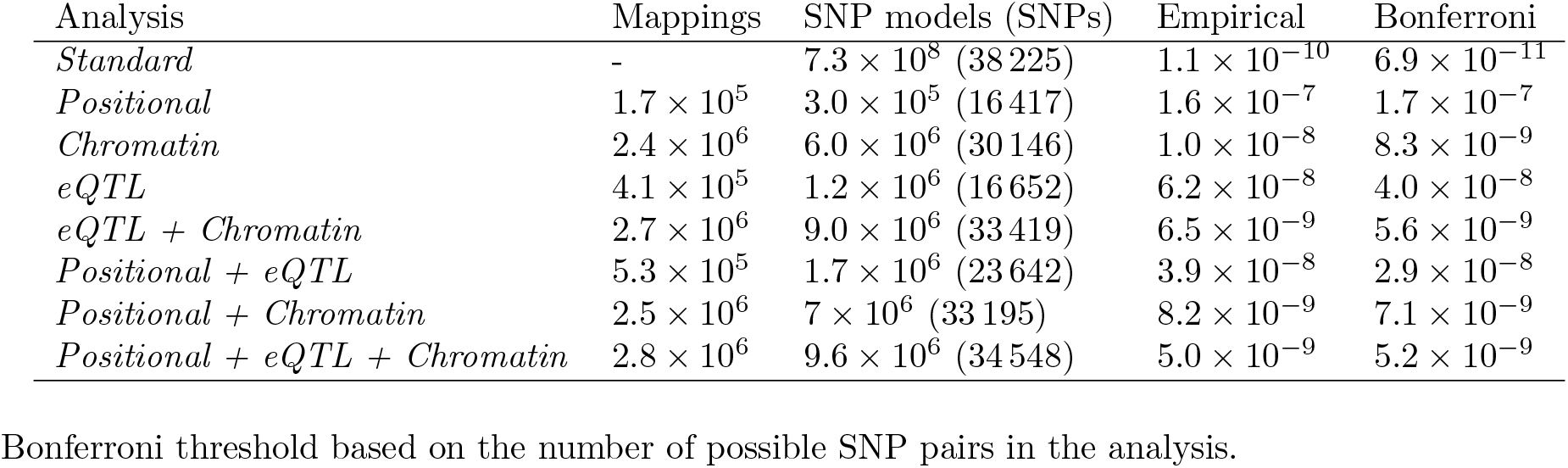
Properties of the different SNP-gene mappings and the filtered datasets. We show the empirical threshold of significance obtained through permutation, and the corresponding Bonferroni threshold for comparison.

| Analysis | Mappings | SNP models (SNPs) | Empirical | Bonferroni |
| --- | --- | --- | --- | --- |
| *Standard* | - | $7.3 \times 10^8$ (38 225) | $1.1 \times 10^{-10}$ | $6.9 \times 10^{-11}$ |
| *Positional* | $1.7 \times 10^5$ | $3.0 \times 10^5$ (16 417) | $1.6 \times 10^{-7}$ | $1.7 \times 10^{-7}$ |
| *Chromatin* | $2.4 \times 10^6$ | $6.0 \times 10^6$ (30 146) | $1.0 \times 10^{-8}$ | $8.3 \times 10^{-9}$ |
| *eQTL* | $4.1 \times 10^5$ | $1.2 \times 10^6$ (16 652) | $6.2 \times 10^{-8}$ | $4.0 \times 10^{-8}$ |
| *eQTL + Chromatin* | $2.7 \times 10^6$ | $9.0 \times 10^6$ (33 419) | $6.5 \times 10^{-9}$ | $5.6 \times 10^{-9}$ |
| *Positional + eQTL* | $5.3 \times 10^5$ | $1.7 \times 10^6$ (23 642) | $3.9 \times 10^{-8}$ | $2.9 \times 10^{-8}$ |
| *Positional + Chromatin* | $2.5 \times 10^6$ | $7 \times 10^6$ (33 195) | $8.2 \times 10^{-9}$ | $7.1 \times 10^{-9}$ |
| *Positional + eQTL + Chromatin* | $2.8 \times 10^6$ | $9.6 \times 10^6$ (34 548) | $5.0 \times 10^{-9}$ | $5.2 \times 10^{-9}$ |

Bonferroni threshold based on the number of possible SNP pairs in the analysis.

### 3.6 Biofilter boosts discovery of interpretable hypotheses

Searching for epistatic interactions exclusively across edges of the Biofilter network greatly reduces the number of tests. Yet, this gain in statistical power might not lead to greater discoveries as it potentially disregards new interactions absent from databases. Hence, we tested whether exhaustively searching for epistasis on the datasets not reduced for Biofilter models but using each mapping, led to similar results. At the SNP level (Fig 5A, upper panel), only a small proportion of the significant interactions were still detected when the network was not used. Strikingly, that difference got smaller at the gene level (Fig 5A, lower panel). This suggests that the significant SNP models, even if fewer in number, are strong enough to lead to the detection of the gene models.

**Fig. 5:**
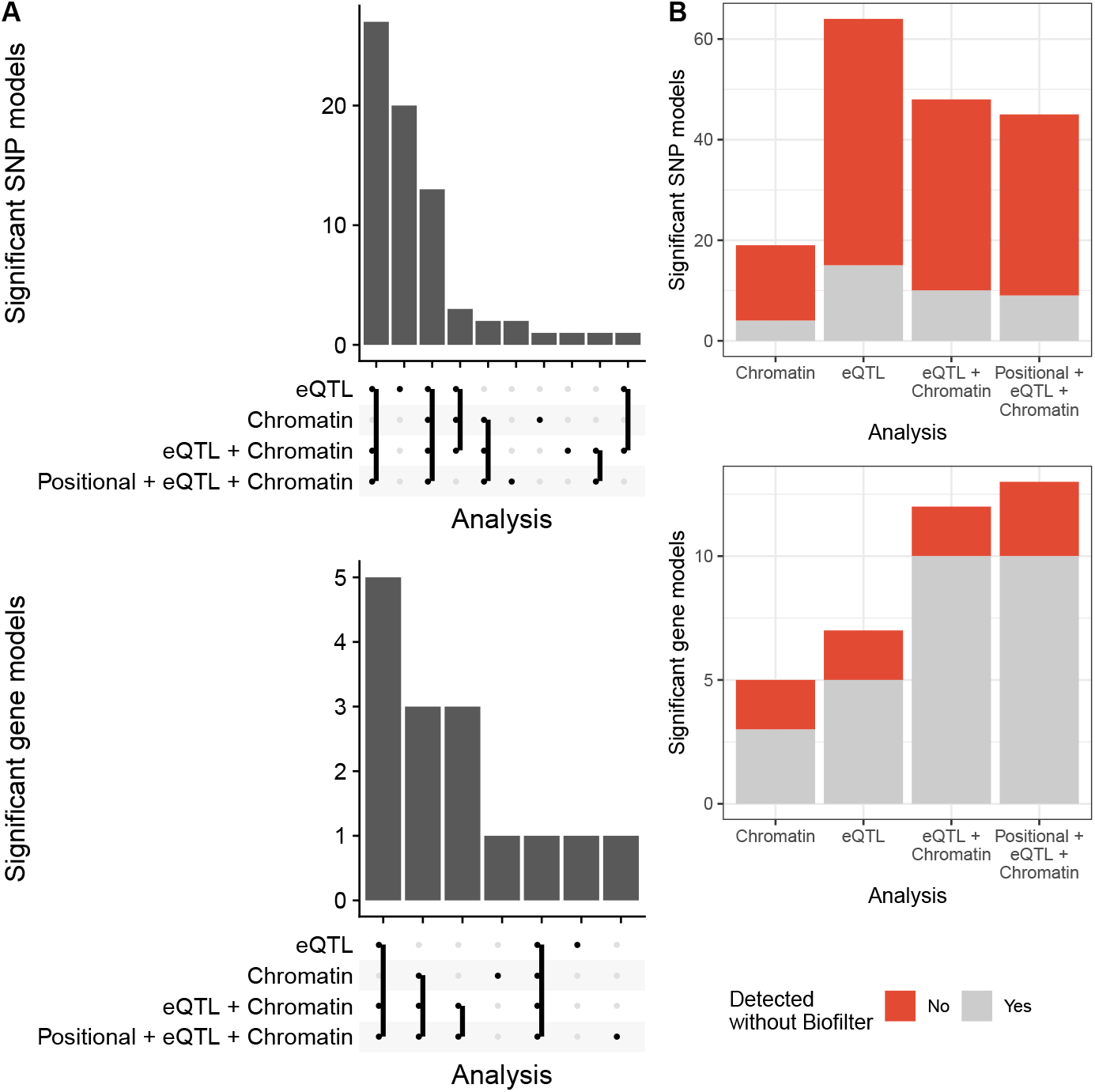
Comparison of the proposed analysis with relaxation of filters at different stages. **(A)** Impact of focusing on one SNP-gene mapping at a time, or at multiple at once. Overlap between the significant interactions detected in the different analyses. SNP interactions on top; gene interactions on the bottom. **(B)** Impact of focusing on interactions mappable to Biofilter interactions. Proportion of significant interactions that were detected using with and without filtering by SNPs mappable to Biofilter interactions. SNP interactions on top; gene interactions on bottom.

In a similar vein, we studied the overlap between the significant models detected in the different analyses. Including more SNP-gene mappings in the analysis was mostly beneficial with respect to considering one mapping at a time, since both at the gene and at the SNP level, the significant interactions in *Positional + eQTL + Chromatin* highly overlapped with the other analyses (Fig 5B). Nonetheless, a few interactions were also missed in this joint analysis, in particular 20 significant SNP models detected in the *eQTL* analysis.

### 3.7 *Chromatin* and *Standard* analyses partially replicate previous studies on IBD

In the past, several genetic studies have investigated epistasis on IBD [14, 25, 26, 30, 33, 45]. We compared them to our results at the gene level, the minimal functional unit at which we expect genetic studies to converge. Several epistatic alterations have been reported involving interleukins [14, 26, 30]. Also our *Standard* analyses resulted in interactions involving three interleukins (*IL-19*, *IL-10* and *IL-23*), although interacting with different genes than in the aforementioned studies. *Positional + eQTL + Chromatin* recovered five interleukins (*IL-4*, *IL-5*, *IL-13*, *IL-19*, *IL-20*). In addition, Lin et al. [25] detected interactions involving *NOD2*, with both *IL-23R* and other genes. Our *Standard* analysis highlighted two potentially new epistasic interactions involving *NOD2*. Discoveries in the proposed protocol are guided by plausible biological interactions. Hence, every significant gene model can be traced back to a biological database, therefore producing biological hypotheses. For instance, the gene model *MST1* -*MST1R* is significant in multiple pipelines. Both genes have been linked to IBD, both by themselves [3, 6] and in interaction with other genes [48]. MST1R is a surface receptor of MST1, and, through physical interaction, they play a role in the regulation of inflammation.

### 3.8 Pathway analyses highlight the involvement of the extracellular matrix in IBD

Pathway enrichment analyses of each interaction’s neighborhood allowed us to identify broader biological mechanisms that the significant interaction pairs might be involved in. The *eQTL* analysis produced multiple significant pathways (see Supplementary Table 1), involving the triangle of interactions formed by two genes located in 3p21.31 (*HYAL1*, *HYAL3*) and one in 7q31.32 (*SPAM1*) (Fig 2). The affected pathways were related to the extracellular matrix, and specifically to glycosaminoglycan degradation. Links between the turnover of the extracelular matric and IBD-related inflammation have been reported [35]. More specifically, glycosaminoglycan [40] and hyaluronon [1] degradation products lead to inflammatory response. When restricting attention to pathways of minimum gene size 10 and maximum gene size 500 to avoid imbalances and non-normality, four pathways are removed: cellular response to UV B, hyalurononglucosaminidase activity, hexosaminidase activity and CS/DS degradation. The *Chromatin* mapping and the *Standard* pipeline did not produce significant pathways. The *Positional + eQTL + Chromatin* analysis produced 71 significant pathways (Supplementary Table 2), involving the neighborhoods *HYAL3*, *HYAL1*, *HYAL2* and *PLA2G2E*, *PLA2G5*, *PLA2G2C*.

### 3.9 The proposed pipeline increases robustness

We studied whether our proposed pipeline led to more robust results. For that purpose, we ran the whole protocol again on a random subset of the data containing 80% of the samples. We repeated this experiment 10 times for each SNP-gene mapping. In each subset, 49% of the individuals were cases, respecting the initial proportion of cases and controls of the entire dataset. Conservatively, we used the same SNP and gene significance thresholds as for the corresponding entire dataset.

The *Standard* pipeline, which does not include Biofilter network-information, produced on average 11.4 significant gene models (standard error (SE) 1.1). With the *eQTL* (respectively *Chromatin*) analysis, we detected on average 5.8 gene pairs (respectively 3.2) with SE 0.1 (respectively 0.4). With the *Positional + eQTL + Chromatin* mappings process, we detected on average 8.6 gene pairs with SE 1.3.

Fig 6 shows that pipelines including biological knowledge recover more than 60% of the gene pairs detected with the entire cohort, on average, (83% for *eQTL*, 60% for *Chromatin* mapping and 64% for *Positional + eQTL + Chromatin*), whereas without this knowledge (*Standard*), we recover less than 40% of the pairs. Hence, the *Standard* analysis appears to be the less robust in terms of conservation of gene pairs. This shows that filtering does increase robustness at the gene level.

**Fig. 6:**
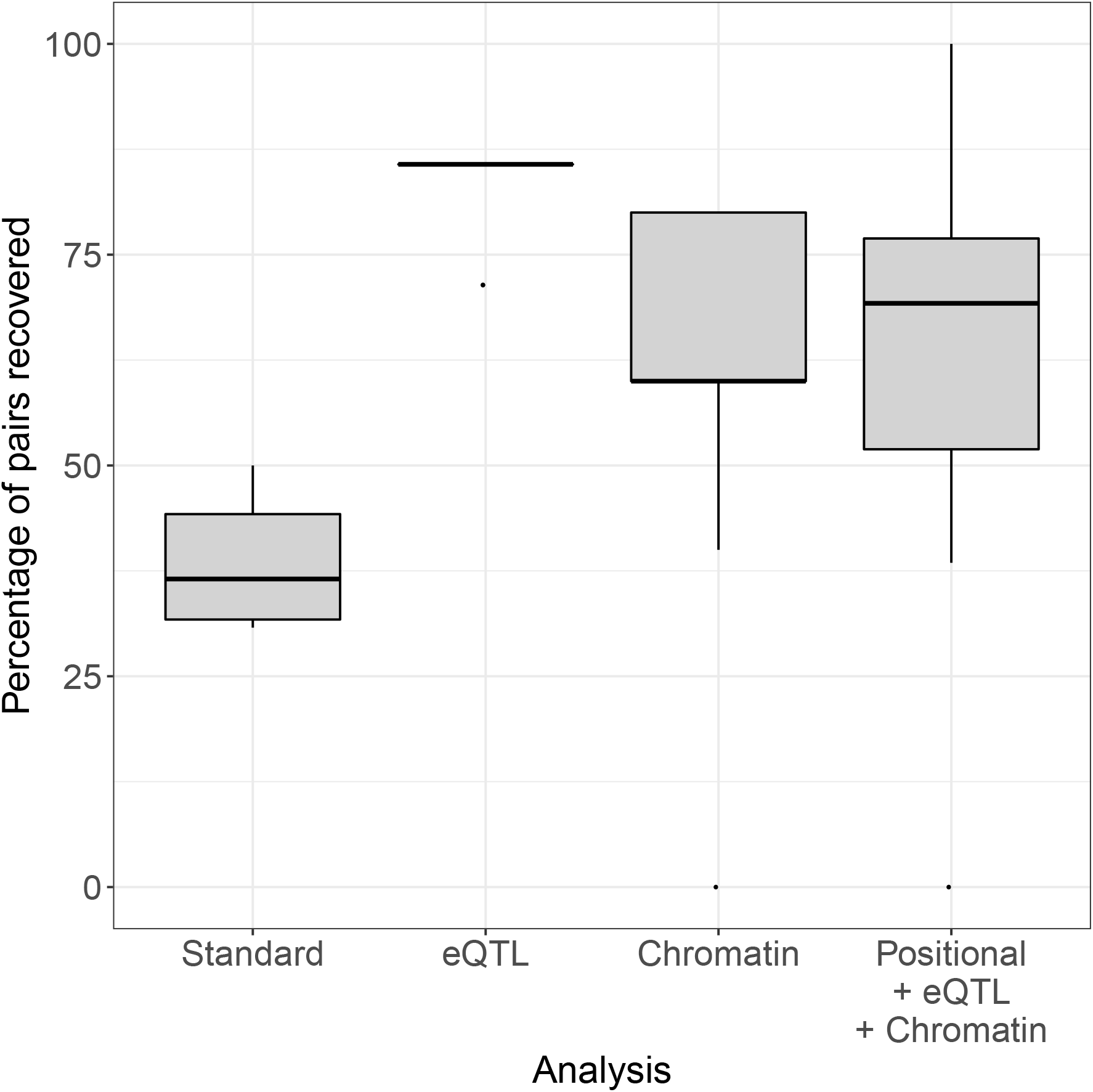
Gene pairs produced within each mapping across ten repetitions using 80% of the data, i.e. percentage of gene pairs detected with the entire dataset that are recovered in the ten subsets with 80% of the individuals.

### 3.10 Tissue-specific mappings do not recover many new interactions

To analyze the impact of tissue-specific mappings, we ran two analyses using exclusively eQTL and Chromatin mappings obtained from relevant tissue types. Specifically, we used mappings obtained from organs and tissues from the nervous and digestive systems (Supplementary Table 3). While the tissue-specific Chromatin analysis did not produce any significant gene pair, the tissue-specific eQTL analysis produced 4 significant gene pairs (Supplementary Fig 4). Nonetheless, only one is novel with regards to the (organism-wide) *eQTL* analysis: *IL18RAP-IL18R1*.

## 4 Discussion

In this article we proposed a new protocol for epistasis detection, based on a variety of functional filtering strategies, and studied its application to GWAS data for Inflammatory Bowel Disease. The protocol included several components to control for type I error, hereby strengthening our belief in the discovered genetic interactions.

A common theme in the interpretation of epistasis results consists on linking the associated variants to an altered gene function. In this article, we considered 3 different such SNP-gene mappings. Notably, the number of SNP-gene correspondences provided by each mapping differed by orders of magnitude. Moreover, the different mappings unevenly described genes; for instance, genes that had most SNPs mapped by using a chromatin contact map, had comparatively few eQTL SNPs. This motivated combining multiple mappings into an analysis (*Positional + eQTL + Chromatin*) in order to combine different perspectives of the epistasis process. For the most part, this complementary approach improved the analysis, by recovering most of the interactions significant in the analyses that used one mapping at a time. Importantly, our results display the benefits of going beyond one single SNP-gene mapping (often, genomic position) to interpret epistasis results. To our surprise, tissue-specific analyses using exclusively eQTL and Chromatin mappings from tissues related to IBD resulted in less significant gene pairs. Despite this setback, we believe that more targeted analyses (e.g. using only interactions from open chromatin in relevant cell-types) might lead to novel discoveries.

Restricting the tested interactions to functionally plausible pairs of genes and SNPs joins two faces of epistasis: searching for statistical epistasis, yet exclusively on plausible candidates for biological epistasis. This has several advantages. First, a more targeted input dataset reduces the number of tests and, in consequence, the multiple testing burden. In contrast, the high dimensionality of GWAIS data requires a much more stringent multiple testing correction and limits the detection of epistasis with low effect sizes. Adopting one of the proposed analyses may reduce the number of SNP interactions to test by more than half (Fig 1). Yet, the *Standard* analysis, which does not use Biofilter, produced the most significant gene models. Second, the proposed protocol addresses the robustness issues widespread in GWAIS by producing results that are consistent at the gene and pathway levels (Fig 5). Indeed, we observed an increased analytic robustness when using Biofilter gene models, in line with previous reports [4]. In particular, *eQTL* and *Chromatin* mappings increased said robustness. Third, restricting the search for epistasis to biologically plausible interactions yields results that are biologically interpretable and strikingly different from the ones obtained without using functional filtering (Fig 2). Not surprisingly, different mappings also provided very different interaction signals and give resolution of information on different genes. In particular, we corroborated that the significant gene models from different functional filters were relevant to the biology of IBD. This was especially true for the *Chromatin* analysis (but also the *eQTL* analysis), giving rise to interactions with seemingly meaningful biological underpinnings, and stressing the relevance of regulatory variants in susceptibility to IBD. In contrast, the *Standard* analysis detected multiple interactions that were hard to interpret. For instance, several interactions involved RNA genes of unknown function (e.g. *LOC101927272* or *LINC02178*).

Remarkably, while the *Standard* analysis produced rich results, the *Positional* analysis did not lead to any significant SNP model. They both use genomic position to map SNPs to genes, but *Positional* is restricted to gene models in Biofilter. We note that the *Positional* analysis does not coincide with how Biofilter is typically used on GWAS data for epistasis detection. The latter involves pooling all SNPs that are mapped to genes which occur in Biofilter proposed gene interaction models, and subsequently exhaustive screening those SNPs for pairwise interactions. These pairs may also involve gene pairs that were not highlighted by Biofilter, in contrast to our *Positional* analysis. We evaluated the impact of Biofilter on the final results. No significant SNP interaction were detected in the *Positional* analysis. In the analysis without biofiltering (dataset reduced to mappable SNPs using genomic proximity, but not reduced to Biofilter gene pairs), 62 pairs were significant. Also, on the 86 SNP interactions that passed the experimental threshold in the *Standard* analysis (dataset not reduced to mappable using genomic proximity, nor Biofilter gene pairs), only 57 are mappable to gene interactions using genomic proximity. Hence, 66% of significant SNP pairs are mappable via genomic proximity in the *Standard* analysis.

An important component of our protocol is the conversion of SNP-based tests to gene-based tests. The most popular approach consists in aggregating SNP-level P-values into gene-level statistics, which can be done in different ways (see Ma et al. [28] for some early examples, and Vsevolozhskaya et al. [46] for recent developments). Here, we developed a generic approach that exploits a permutation strategy to define a P-value cutoff for SNP interactions, at a FWER of 5%, and then we followed the original implementation of the adaptive TPM (ATPM) to accommodate several truncation thresholds at once [39] while taking permutations instead of bootstrap as in Yu et al. [51]. The two algorithms are very similar, but we favored the TPM over the rank truncated product method of Yu et al. [51] that employs the product of the L most significant P-values, because the TPM only requires P-values smaller than a specified threshold, which is in line with the output of PLINK epistasis detection and saves storage space. Following both protocols and the recommendation of Becker and Knapp [2] we included measures derived from observed data in computing statistics under the null.

Remarkably, our proposed procedure keeps type I error under control, without additional corrections for multiple testing at the gene model. We hypothesize that this stems from two reasons. First, we apply a stringent correction for multiple testing at the SNP level. Second, when moving from SNP model significance to gene model significance, the ATPM only considers gene models that map to at least one significant SNP model. However, alternative strategies could have been considered. For instance, not restricting ourselves to significant SNP models, hence conducting ATPM on all gene models. This could have led to increased discovery, in cases where the SNP models mapped to a gene tend to be low, albeit non-significant. However, it may also lead to an increased type I error. Accounting for that would require a multiple test correction at the gene level. In turn, such correction would be difficult since the dependency between the tests is unknown. Additionally, in common multiple test correction procedures this would require a much higher number of permutations, in order to obtain the necessary numerical precision.

How to best perform a pathway analysis of epistasis results is understudied. Often, all genes belonging to any significant gene pair are simply pooled together into a joint enrichment analysis. This approach discards the gene-gene interaction information that was, indeed, the object of analysis. Hence, in our procedure we adapted the *Network neighborhood search* protocol from Yip et al. [50], which considers the topology of the network using the shortest paths between the studied genes. It should be noted that we only used the topology to derive a neighborhood for each significant gene pair; then, we discarded the edge information. Yet, there are several directions for improvement. One is to exploit the topology of the epistasis network beyond the creation of a neighborhood. Another one is to take into account the gene size (or the number of SNPs per gene), for instance by performing a weighted version of the statistical test. Jia et al. [20] suggested a method for gene set enrichment analysis of GWAS data, adjusting the gene length bias or the number of SNP per gene. In our data, we did observe a link between the significance of the gene models and the number of SNPs mapped to the gene. For instance, in the *eQTL* analysis, the only one producing significant pathways, the median number of SNPs per genes is 385 among genes in significant pairs, versus 3 SNPs/gene genome-wide.

Several protocol changes may impact final results. As reported elsewhere [4], these changes or choices include the modelling framework (parametric, non-parametric, semi-parametric), encoding of the genetic markers, as well as LD handling. In this work, we used an additive encoding scheme (0, 1, 2 indicating the number of copies of the minor SNP allele), a popular choice in part because of its computational efficiency. However, this encoding schemes has been reported to tend to increase false positives (for instance [44]). This observation is based on type I error studies with data generated under the null hypothesis of no pairwise genetic interactions but in the presence of main effects (see for instance [21]). Here, we investigated the type I error control of our protocols under a general null hypothesis of no genetic associations with the trait (no interactions and no main effects) and established adequate control. As a consequence, this does not guarantee that our generated SNP interaction results were not overly-optimistic. To this end, we adjusted SNP-level epistasis P-values for main effects as comprised in a polygenic risk score. Not only does such a post-analysis adjustment via conditional regression reduce over-optimism due to inadequate control for lower-order effects, thus addressing phantom epistasis [10], but it may also occasionally highlight the masking of SNP interactions (as was shown in Fig 4B - *eQTL*). More work is needed to investigate the impact on gene-level interaction results, derived accordingly. For convenience, we used the regression framework to identify SNP interactions and relied on earlier recommendations regarding LD handling [18].

Our protocols are built on output from Biofilter, that can be presented as a co-functional gene network. One of the motivations was its proven ability to highlight meaningful interactions in a narrower alternative hypothesis space, at the expense of leaving parts of the interaction search space unexplored. The database that Biofilter built contained 37 266 interactions. This is notably smaller than other gene interaction databases, like HINT [9], 173 797 interactions), or STRING [42], 11 759 455 interactions). Testing gene interactions with other (combinations of) biological interaction networks was beyond the scope of this paper. Furthermore, Biofilter analysis or exhaustive screening may lead to non-overlapping results. An example within a regression context is given in [4].

## 5 Potential implications

In this study we presented a protocol to enhance the interpretation of epistasis screening from GWAS. It includes gene-level epistasis discoveries with type I error under control, as well as a network-guided pathway analysis. Moreover, it improves the robustness of the results. While SNP pairs from a GWAIS study rarely replicate in other cohorts and arrays, results at the gene and pathway level are more likely to be reproducible. This can be achieved directly by applying the proposed protocol, or by testing SNP models in a cohort obtained from the gene pairs and pathways significant in other studies, via the SNP-gene mapping of interest. Aggregating SNP-level results into gene-level epistasis is challenging, but allows to include information from biological interaction databases. Based on that, we conducted multiple analyses that use different sources of prior biological knowledge about SNP-to-gene relationships and gene interaction models, as well as rigorous statistical approaches to assess significance. Each of them offers a different, albeit complementary view of the disease, which leads to additional insights.

Their application to GWAS data for inflammatory bowel disease highlighted the potential of our strategy, including network-guided pathway analysis, as it recovered known aspects of IBD while capturing relevant and previously unreported features of its genetic architecture. These strategies will contribute to identify gene-level interactions from SNP data for complex diseases, and to enhance our belief in these findings.

## 6 Methods

### 6.1 Gene interaction detection procedure

As we describe in more detail below, we applied different functional filters to the available data. These filters use plausible interactions between genes, and three different ways of mapping SNPs to those genes, and hence, to these interactions. These three mappings exploit different degrees of biological knowledge to map SNPs to genes, referred to as *Positional*, *eQTL* and *Chromatin*. For each of the three SNP-to-gene mappings, we only analyzed the pairs of SNPs corresponding to a gene pair with prior evidence for interaction. Across this article, we compared our findings in these scenarios to a *Standard* scenario. In this case, we exhaustively search for epistasis between all 38 225 SNPs that passed quality control (Table 4). We mapped the resulting significant SNP interactions to potential gene interactions using the positional mapping. An overview of the entire pipeline is presented in Fig 7.

**Fig. 7:**
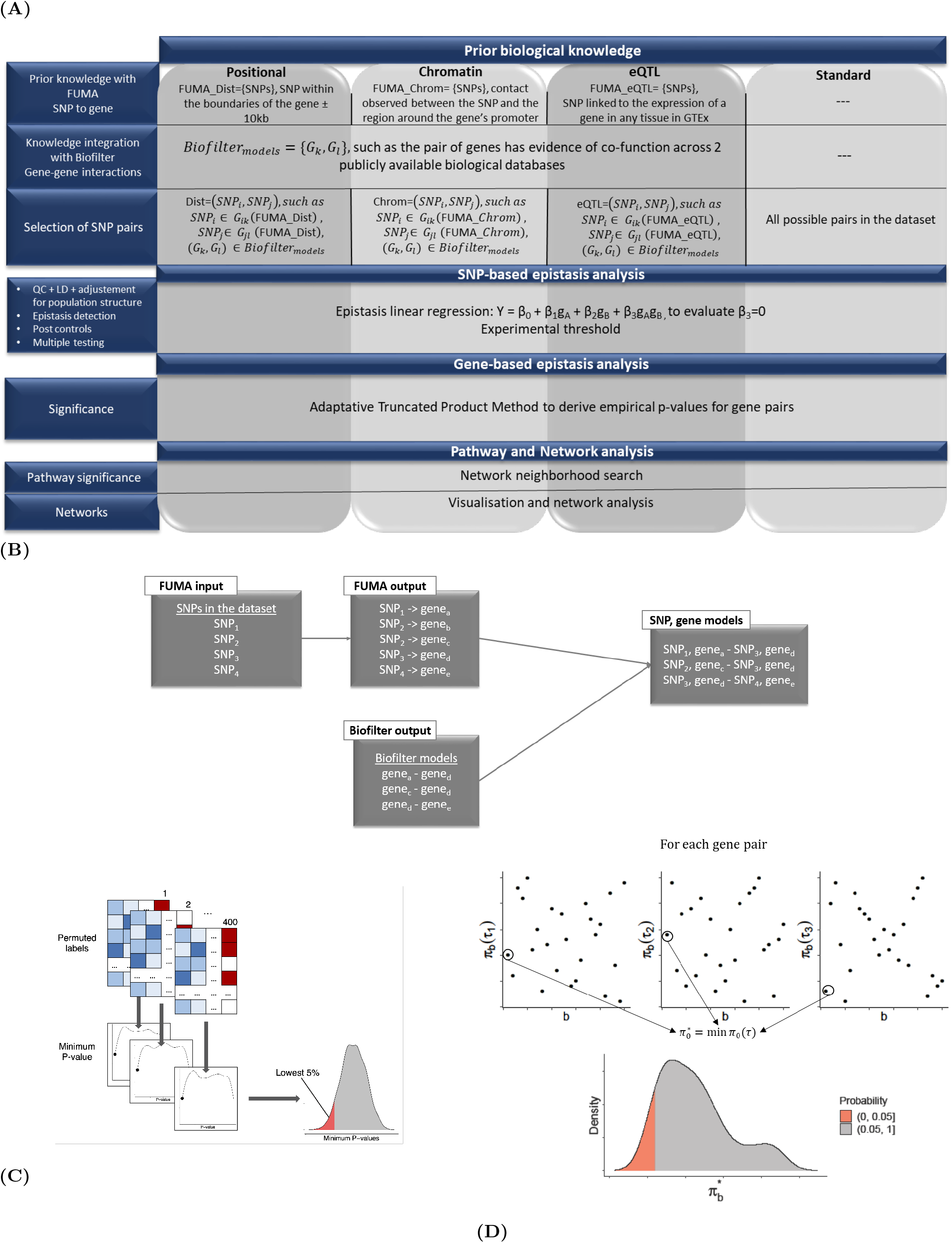
**(A)** Overview of the investigated gene-gene interaction detection protocols, described in *Gene interaction detection procedure*. **(B)** Summary of the procedure to obtain SNP and gene models using FUMA and Biofilter, described in *Co-function gene and SNP networks*. **(C)** Permutation procedure to obtain the SNP model P-value threshold, described in *SNP-level epistasis detection and multiple testing correction*. **(D)** Overview of the adaptive truncated product methodology, described in *From SNP-level to gene-level epistasis*.

#### 6.1.1 From gene models to SNP models

Although the unit of analysis in GWAIS is the SNP, biological interactions are often characterized at the gene level. Hence, we mapped all SNPs in the dataset to genes using FUMA [47], a post-GWAS annotation tool. We created an artificial input where every SNP is significant in order to perform such mapping on all the SNPs. We performed three SNP-gene mappings using FUMA’s SNP2GENE: positional, eQTL and 3D chromatin interaction (Table 4). In the *Positional* mapping, we mapped a SNP to a gene when the genomic coordinates of former was within the boundaries of the latter ±10 kb. The *eQTL* mapping uses eQTLs obtained from GTEx [16]. We mapped an eQTL SNP to its target gene when the association P-value was significant in any tissue (FDR < 0.05). Lastly, in the *Chromatin* mapping, we mapped a SNP to a gene when a contact had been observed between the former and the region around the latter’s promoter in the 3D structure of the genome (250 bp upstream and 500 bp downstream from the transcription start site) in any of the Hi-C datasets included in FUMA (FDR < 10^−6^). This mapping might contain new, undiscovered, regulatory variants which, as for SNPs obtained through eQTL mapping, regulate the expression of a gene.

#### 6.1.2 Co-function gene and SNP networks

We used Biofilter 2.4 [34] to obtain candidate gene pairs to investigate for epistasis evidence. Biofilter generates pairs of genes susceptible to interact (*gene models*) with evidence of co-function across multiple publicly available biological databases. It includes genomic locations of SNPs and genes, as well as known relationships among genes and proteins such as interaction pairs, pathways and ontological categories, but does not use trait information. As per Biofilter’s default, we used gene models supported by evidence in at least 2 databases. Additionally, we removed self-interactions, as detection of within-gene epistasis requires special considerations and is beyond the scope of this paper.

Given this set of gene models, and three different ways of obtaining *SNP models* from it, we removed all the SNPs that did not participate in any SNP model. Subsequently, we created eight datasets. In one dataset no filter was applied (*Standard* analysis), i.e. no Biofiltering nor any SNP-to-gene mapping. Hence, the original SNP set was used. We also created one dataset exclusively for each SNP-to-gene mapping (*Positional*, *eQTL* and *Chromatin*). Lastly, we created four datasets using joint mappings: one with all the mappings (*Positional + eQTL + Chromatin*); and three with only two of them (*eQTL + Chromatin*, *Positional + eQTL* and *Positional + Chromatin*).

We discarded SNP models involving rare variants (MAF < 5%) or in Hardy–Weinberg equilibrium (P-value < 0.001). Regardless, all risk SNPs described in Liu et al. [27] were included, even when the aforementioned epistasis quality controls criteria did not hold up. Then, when the two SNPs of a *SNP model* were located in the HLA region, we discarded the pair, as it is difficult to differentiate between main and non-additive effects in this region [43]. Lastly, we discarded models where the SNPs were in linkage equilibrium (*r*^2^ > 0.75), as motivated in Gusareva and Van Steen [18].

#### 6.1.3 SNP-level epistasis detection and multiple testing correction

We used PLINK 1.9 to detect epistasis through a linear regression on the population structure adjusted phenotypes with the option `--epistasis`:

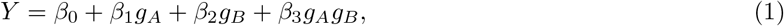

where *g*_*A*_ and *g*_*B*_ are the genotypes under additive encoding for SNPs A and B respectively, *Y* is the adjusted phenotype, and *β*_0_, *β*_1_, *β*_2_, and *β*_3_ are the regression coefficients. PLINK performs a statistical test to evaluate whether *β*_3_ ≠ 0. It only returns SNP pairs with a P-value lower than a specified threshold. We used the default 0.0001. Only SNP models were considered, apart from the *Standard* approach.

To correctly account for multiple testing, the P-value threshold of significance had to be dataset-dependent as the number of tested SNP pairs changed from dataset to dataset. We obtained these thresholds through permutations as in Hemani et al. [19] (Fig 7). More specifically, for each dataset, we permuted the phenotypes 400 times and fitted the aforementioned regression-based association model. This produced a null distribution of the extreme P-values for this number of tests given the LD structure in the data. For each dataset, we took the most extreme P-value from each of the 400 permutations and set the threshold for 5% family-wise error rate (FWER) to be the 5% percentile of these most extreme P-values. Posterior experiments showed that a higher number of permutations, 1 000, barely changed the empirical threshold (data not shown). Hence, 400 was a sufficient number of permutations to obtain an adequate threshold.

#### 6.1.4 From SNP-level to gene-level epistasis

Our next step was to use significant SNP interactions to identify significant gene interactions, which requires combining the P-values of all SNP pairs mapped to the same gene pair. Suppose that SNP interaction tests have been conducted for *N* individual hypotheses *H*_0*i*_, *i* = 1, 2, …, *N*, for example, *N* SNP models mapped to the same gene model. We tested the joint null hypothesis 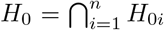 at significance level *α* versus the combined alternative hypothesis *H*_1_: at least one of *H*_0*i*_ is false. To do so, we considered all SNP pairs mapped to the same gene pair as a set of tests with the same global null hypothesis, and applied the Adaptive Truncated Product Method (ATPM) [39] (Fig 7).

ATPM is an adaptive variant of the Truncated Product Method (TPM) of Zaykin et al. [52], which uses as a statistic the product of the P-values smaller than some pre-specified threshold (here, significant SNP interactions) tests. More specifically, given a truncation point *τ* and a number *N* of significant SNP interactions, this test statistic is given as 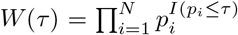 where *I*(·) is the indicator function. TPM is interesting in our context because it does not require P-values for all SNP pairs but only for the most strongly associated ones.

The distribution of *W* (τ) under the null hypothesis is unknown when the individual tests are not independent, which is clearly the case here, but an empirical P-value 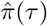 can be estimated through permutations. Because the choice of *τ* is arbitrary, the adaptive version of TPM (ATPM) explores several values of *τ* and selects 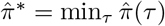. The distribution of 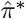 under the null hypothesis can again be determined through permutations [13].

In our procedure, which is detailed below for a given gene pair, we used *B* = 999 permutations and *τ* ∈ {0.001, 0.01, 0.05}. Remarkably, and following the suggestion of Becker and Knapp [2], the null distribution includes both the statistic from the observed dataset, and from the 999 permutations.

1. For each SNP model *i* = 1, *…, N* mapped to the gene pair, compute its P-values *p_i,b_* in the original dataset (*b* = 0) and for each of the *B* = 999 permutations (*b* = 1, *…, B*).
2. For each value of *τ* and *b*, compute the test statistic *W* (*τ*).
3. For each value of *τ* and *b*, estimate the P-value : 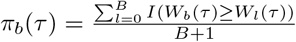.
4. For each value of *b*, compute 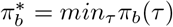.
5. Estimate the P-value of the gene model as 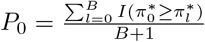.
6. Reject the global null hypothesis if *P*_0_ ≤ *α* = 0.05.

### 6.2 Studying the impact of confounding main effects

The SNPs from some detected interactions were near SNPs with main effects. To assess the impact on the results, we studied the difference between *β*_3_ in Eq. 1 and in the following model:

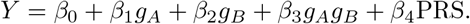

PRS is the polygenic risk score (PRS) computed for the sample. We expect the PRS to capture the variance explained by main all effects.

We computed the PRS with PRSice-2 [7], using the default options. Since it requires GWAS summary statistics, we used PLINK `--assoc` to compute the association of each SNP in the original dataset (130 071 SNPs and 66 280 individuals, with the trait adjusted for PCs). Since the adjusted phenotype is quantitative, PLINK computes the linear regression coefficients and assesses their significance using the Wald test. PRSice performs clumping to remove SNPs that are in LD with each other. The *r*^2^ values computed by PRSice are based on maximum likelihood haplotype frequency estimates. From the 130 071 initial variants, 28 389 variants remained after clumping (`--clump-kb 250kb`, `--clump-p 1`, `--clump-r2 0.1`). We used the average effect size method to calculate the PRS, with high-resolution scoring.

### 6.3 Pathway analysis

A pathway enrichment analysis on the neighborhood of a significant gene model can inform about the broader framework in which gene epistasis occurs. To define such neighborhoods, we adapted the network neighborhood search protocol from Yip et al. [50]. We computed the neighborhood of two genes as the list of all genes that (1) participate in any of the shortest paths between the two studied genes in the Biofilter network, once the direct link between them is removed; and (2) are also involved in a significant interaction with at least one other gene on these paths. We restricted our attention to neighborhoods containing at least 3 genes, including the 2 from the considered gene model. For each of these, we conducted a gene set enrichment analysis in all human gene sets from the Molecular Signature Database (MSigDB version 7) [24, 41]. We performed the enrichment analysis using a hypergeometric test, which compares the obtained overlap between two sets to the expected overlap from taking equally-sized random sets from the universe of genes. We favored the hypergeometric test over the chi-square test used in Yip et al. [50] because the sample sizes of the neighborhoods were small and because chi-square is an approximation whereas the hypergeometric test is an exact test. The universe set was analysis dependant. It contained all genes in an annotated pathway and that can be mapped via genomic proximity to a SNP of the dataset for the *Standard* analysis, and genes present in Biofilter gene models, in an annotated pathway and that can be mapped via the appropriate SNP to gene mapping to a SNP of the dataset for the other analysis. Finally, pathways were said to be significant when the corresponding test P-value was lower than the Bonferroni threshold (0.05*/*(# pathways×# tested gene neighborhoods)), with *pathways* corresponding to pathways containing at least one gene of the neighborhood.

## 7 Availability of source code and requirements

- Project name: `network epistasis.nf`
- Project home page: https://github.com/hclimente/gwas-tools
- Operating system(s): Platform independent
- Programming language: nextflow
- Other requirements: Bash, nextflow, PLINK 1.9 and R 4.0 or higher, with the following packages: clusterProfiler, data.table, igraph, msigdbr, snpStats, tidyverse.
- License: GNU GPL v3.0

Additionally, the code necessary to reproduce this article’s results and analyses is available on GitHub at https://github.com/DianeDuroux/BiologicalEpistasis.

## 8 Data Availability

The data set supporting the results of this article is available upon request from the International Inflammatory Bowel Disease Genetics Consortium (https://www.ibdgenetics.org/). GWAS summary statistics are publicly available.

## 9 Declarations

### 9.1

#### List of abbreviations

ATPM: adaptive truncated product method
eQTL: expression quantitative trait loci
FWER: family-wise error rate
GWAS: genome-wide association study
GWAIS: genome-wide association interaction study
IBD: inflammatory bowel disease
LD: linkage disequilibrium
SNP: single-nucleotide polymorphism
TPM: truncated product method

### 9.2 Ethical Approval

Not applicable.

### 9.3 Consent for publication

Not applicable.

### 9.6 Authors’ contributions

Conceptualization: D.D., H.C.G., C-A.A., K.V.S.; data curation: D.D., H.C.G.; formal Analysis: D.D., H.C.G.; funding acquisition: C-A.A., K.V.S.; investigation: D.D., H.C.G.; methodology: D.D., K.V.S.; project administration: C-A.A., K.V.S.; resources: C-A.A., K.V.S.; software: D.D., H.C.G.; supervision: C-A.A., K.V.S.; validation: D.D., H.C.G.; visualization: D.D., H.C.G.; writing – original draft: D.D., H.C.G.; writing – review & editing: D.D., H.C.G.

## 10 Acknowledgments

We thank the International IBD Genetics Consortium for data collection and processing and for interesting discussions.

## References

[1] S. Albeiroti, A. Soroosh, and C. A. de la Motte. Hyaluronan’s role in fibrosis: a pathogenic factor or a passive player? BioMed research international, 2015, 2015.

[2] T. Becker and M. Knapp. A powerful strategy to account for multiple testing in the context of haplotype analysis. The American Journal of Human Genetics, 75(4):561–570, 2004.

[3] J. B. Beckly, L. Hancock, A. Geremia, F. J. Cummings, A. Morris, R. Cooney, S. Pathan, C. Guo, and D. P. Jewell. Two-stage candidate gene study of chromosome 3p demonstrates an association between nonsynonymous variants in the MST1r gene and crohn’s disease. Inflammatory Bowel Diseases, 14(4): 500–507, Apr. 2008. doi: 10.1002/ibd.20365. URL https://doi.org/10.1002/ibd.20365.

[4] K. Bessonov, E. S. Gusareva, and K. Van Steen. A cautionary note on the impact of protocol changes for genome-wide association snp× snp interaction studies: an example on ankylosing spondylitis. Human genetics, 134(7):761–773, 2015.

[5] A. Buniello, J. A. MacArthur, M. Cerezo, L. W. Harris, J. Hayhurst, C. Malangone, A. McMahon, J. Morales, E. Mountjoy, E. Sollis, D. Suveges, O. Vrousgou, P. L. Whetzel, R. Amode, J. A. Guillen, H. S. Riat, S. J. Trevanion, P. Hall, H. Junkins, P. Flicek, T. Burdett, L. A. Hindorff, F. Cunningham, and H. Parkinson. The NHGRI-EBI GWAS Catalog of published genome-wide association studies, targeted arrays and summary statistics 2019. Nucleic Acids Research, 47(D1):D1005–D1012, Jan. 2019. ISSN 0305-1048, 1362-4962. doi: 10.1093/nar/gky1120. URL https://academic.oup.com/nar/article/47/D1/D1005/5184712.00092.

[6] C. P. R. Burton, D. G. Clayton, L. R. Cardon, N. Craddock, P. Deloukas, A. Duncanson, D. P. Kwiatkowski, M. I. McCarthy, W. H. Ouwehand, N. J. Samani, J. A. Todd, P. Donnelly, J. C. Barrett, P. R. Burton, D. Davison, P. Donnelly, D. Easton, D. Evans, H.-T. Leung, J. L. Marchini, A. P. Morris, C. C. A. Spencer, M. D. Tobin, L. R. Cardon, D. G. Clayton, A. P. Attwood, J. P. Boorman, B. Cant, U. Everson, J. M. Hussey, J. D. Jolley, A. S. Knight, K. Koch, E. Meech, S. Nutland, C. V. Prowse, H. E. Stevens, N. C. Taylor, G. R. Walters, N. M. Walker, N. A. Watkins, T. Winzer, J. A. Todd, W. H. Ouwehand, R. W. Jones, W. L. McArdle, S. M. Ring, D. P. Strachan, M. Pembrey, G. Breen, D. St Clair, S. Caesar, K. Gordon-Smith, L. Jones, C. Fraser, E. K. Green, D. Grozeva, M. L. Hamshere, P. A. Holmans, I. R. Jones, G. Kirov, V. Moskvina, I. Nikolov, M. C. O’Donovan, M. J. Owen, N. Craddock, D. A. Collier, A. Elkin, A. Farmer, R. Williamson, P. McGuffin, A. H. Young, I. N. Ferrier, S. G. Ball, A. J. Balmforth, J. H. Barrett, D. T. Bishop, M. M. Iles, A. Maqbool, N. Yuldasheva, A. S. Hall, P. S. Braund, P. R. Burton, R. J. Dixon, M. Mangino, S. Stevens, M. D. Tobin, J. R. Thompson, N. J. Samani, F. Bredin, M. Tremelling, M. Parkes, H. Drummond, C. W. Lees, E. R. Nimmo, J. Satsangi, S. A. Fisher, A. Forbes, C. M. Lewis, C. M. Onnie, N. J. Prescott, J. Sanderson, C. G. Mathew, J. Barbour, M. K. Mohiuddin, C. E. Todhunter, J. C. Mansfield, T. Ahmad, F. R. Cummings, D. P. Jewell, J. Webster, M. J. Brown, D. G. Clayton, G. M. Lathrop, J. Connell, A. Dominiczak, N. J. Samani, C. A. B. Marcano, B. Burke, R. Dobson, J. Gungadoo, K. L. Lee, P. B. Munroe, S. J. Newhouse, A. Onipinla, C. Wallace, M. Xue, M. Caulfield, M. Farrall, A. Barton, T. B. i. R. G. Genomics (BRAGGS), I. N. Bruce, H. Donovan, S. Eyre, P. D. Gilbert, S. L. Hider, A. M. Hinks, S. L. John, C. Potter, A. J. Silman, D. P. M. Symmons, W. Thomson, J. Worthington, D. G. Clayton, D. B. Dunger, S. Nutland, H. E. Stevens, N. M. Walker, B. Widmer, J. A. Todd, T. M. Frayling, R. M. Freathy, H. Lango, J. R. B. Perry, B. M. Shields, M. N. Weedon, A. T. Hattersley, G. A. Hitman, M. Walker, K. S. Elliott, C. J. Groves, C. M. Lindgren, N. W. Rayner, N. J. Timpson, E. Zeggini, M. I. McCarthy, M. Newport, G. Sirugo, E. Lyons, F. Vannberg, A. V. S. Hill, L. A. Bradbury, C. Farrar, J. J. Pointon, P. Wordsworth, M. A. Brown, J. A. Franklyn, J. M. Heward, M. J. Simmonds, S. C. L. Gough, S. Seal, B. C. Susceptibility Collaboration (UK), M. R. Stratton, N. Rahman, M. Ban, A. Goris, S. J. Sawcer, A. Compston, D. Conway, M. Jallow, M. Newport, G. Sirugo, K. A. Rockett, D. P. Kwiatkowski, S. J. Bumpstead, A. Chaney, K. Downes, M. J. R. Ghori, R. Gwilliam, S. E. Hunt, M. Inouye, A. Keniry, E. King, R. McGinnis, S. Potter, R. Ravindrarajah, P. Whittaker, C. Widden, D. Withers, P. Deloukas, H.-T. Leung, S. Nutland, H. E. Stevens, N. M. Walker, J. A. Todd, D. Easton, D. G. Clayton, P. R. Burton, M. D. Tobin, J. C. Barrett, D. Evans, A. P. Morris, L. R. Cardon, N. J. Cardin, D. Davison, T. Ferreira, J. Pereira-Gale, I. B. Hallgrimsdóttir, B. N. Howie, J. L. Marchini, C. C. A. Spencer, Z. Su, Y. Y. Teo, D. Vukcevic, P. Donnelly, D. Bentley, M. A. Brown, L. R. Cardon, M. Caulfield, D. G. Clayton, A. Compston, N. Craddock, P. Deloukas, P. Donnelly, M. Farrall, S. C. L. Gough, A. S. Hall, A. T. Hattersley, A. V. S. Hill, D. P. Kwiatkowski, C. G. Mathew, M. I. McCarthy, W. H. Ouwehand, M. Parkes, M. Pembrey, N. Rahman, N. J. Samani, M. R. Stratton, J. A. Todd, J. Worthington, The Wellcome Trust Case Control Consortium, Management Committee, Data and Analysis Committee, UK Blood Services and University of Cambridge Controls, 1958 Birth Cohort Controls, Bipolar Disorder, Coronary Artery Disease, Crohn’s Disease, Hypertension, Rheumatoid Arthritis, Type 1 Diabetes, Type 2 Diabetes, Tuberculosis, Ankylosing Spondylitis, Autoimmune Thyroid Disease, Breast Cancer, Multiple Sclerosis, Gambian Controls, G. DNA, Data QC and Informatics, Statistics, and Primary Investigators. Genome-wide association study of 14,000 cases of seven common diseases and 3,000 shared controls. Nature, 447(7145):661–678, June 2007. ISSN 1476-4687. doi: 10.1038/nature05911. URL https://doi.org/10.1038/nature05911.00000.

[7] S. W. Choi and P. F. O’Reilly. Prsice-2: Polygenic risk score software for biobank-scale data. Gigascience, 8(7):giz082, 2019.

[8] A. Cortes and M. A. Brown. Promise and pitfalls of the Immunochip. Arthritis Research & Therapy, 13(1): 101, 2010. ISSN 1478-6354. doi: 10.1186/ar3204. URL http://arthritis-research.biomedcentral.com/articles/10.1186/ar3204.00451.

[9] J. Das and H. Yu. HINT: High-quality protein interactomes and their applications in understanding human disease. BMC Systems Biology, 6(1):92, 2012. ISSN 1752-0509. doi: 10.1186/1752-0509-6-92. URL http://bmcsystbiol.biomedcentral.com/articles/10.1186/1752-0509-6-92.00204.

[10] G. de los Campos, D. A. Sorensen, and M. A. Toro. Imperfect linkage disequilibrium generates phantom epistasis (& perils of big data). G3: Genes, Genomes, Genetics, 9(5):1429–1436, Mar. 2019. doi: 10.1534/g3.119.400101. URL https://doi.org/10.1534/g3.119.400101.

[11] D. Ellinghaus, L. Jostins, S. L. Spain, A. Cortes, J. Bethune, B. Han, Y. R. Park, S. Raychaudhuri, J. G. Pouget, M. Hübenthal, et al. Analysis of five chronic inflammatory diseases identifies 27 new associations and highlights disease-specific patterns at shared loci. Nature genetics, 48(5):510, 2016.

[12] D. Ellinghaus, S. L. Spain, A. Cortes, J. Bethune, B. Han, Y. R. Park, S. Raychaudhuri, J. G. Pouget, M. Hübenthal, T. Folseraas, Y. Wang, T. Esko, A. Metspalu, H.-J. Westra, L. Franke, T. H. Pers, R. K. Weersma, V. Collij, M. D’Amato, J. Halfvarson, A. B. Jensen, W. Lieb, F. Degenhardt, A. J. Forstner, A. Hofmann, S. Schreiber, U. Mrowietz, B. D. Juran, K. N. Lazaridis, S. Brunak, A. M. Dale, R. C. Trembath, S. Weidinger, M. Weichenthal, E. Ellinghaus, J. T. Elder, J. N. W. N. Barker, O. A. Andreassen, D. P. McGovern, T. H. Karlsen, J. C. Barrett, M. Parkes, M. A. Brown, and A. Franke. Analysis of five chronic inflammatory diseases identifies 27 new associations and highlights disease-specific patterns at shared loci. Nature Genetics, 48(5):510–518, May 2016. ISSN 1061-4036, 1546-1718. doi: 10.1038/ng.3528.URL http://www.nature.com/articles/ng.3528.00214.

[13] Y. Ge, S. Dudoit, and T. P. Speed. Resampling-based multiple testing for microarray data analysis. Test, 12(1):1–77, 2003.

[14] J. Glas, J. Stallhofer, S. Ripke, M. Wetzke, S. Pfennig, W. Klein, J. T. Epplen, T. Griga, U. Schiemann, M. Lacher, et al. Novel genetic risk markers for ulcerative colitis in the il2/il21 region are in epistasis with il23r and suggest a common genetic background for ulcerative colitis and celiac disease. The American journal of gastroenterology, 104(7):1737, 2009.

[15] H. Gordon, F. Trier Moller, V. Andersen, and M. Harbord. Heritability in inflammatory bowel disease: from the first twin study to genome-wide association studies. Inflammatory bowel diseases, 21(6):1428–1434, 2015.

[16] GTEx Consortium. Genetic effects on gene expression across human tissues. Nature, 550(7675):204–213, Oct. 2017. ISSN 0028-0836, 1476-4687. doi: 10.1038/nature24277. URL http://www.nature.com/articles/nature24277.00708.

[17] A. C. Gumpinger, B. Rieck, D. G. Grimm, International Headache Genetics Consortium, and K. Borgwardt. Network-guided search for genetic heterogeneity between gene pairs. Bioinformatics, page btaa581, June 2020. ISSN 1367-4803, 1460-2059. doi: 10.1093/bioinformatics/btaa581. URL https://academic.oup.com/bioinformatics/advance-article/doi/10.1093/bioinformatics/btaa581/5861532.00000.

[18] E. S. Gusareva and K. Van Steen. Practical aspects of genome-wide association interaction analysis. Human Genetics, 133(11):1343–1358, Nov. 2014. ISSN 0340-6717, 1432-1203. doi: 10.1007/s00439-014-1480-y. URL http://link.springer.com/10.1007/s00439-014-1480-y.00015.

[19] G. Hemani, K. Shakhbazov, H.-J. Westra, T. Esko, A. K. Henders, A. F. McRae, J. Yang, G. Gibson, N. G. Martin, A. Metspalu, L. Franke, G. W. Montgomery, P. M. Visscher, and J. E. Powell. Detection and replication of epistasis influencing transcription in humans. Nature, 508(7495):249–253, Apr. 2014. ISSN 0028-0836, 1476-4687. doi: 10.1038/nature13005. URL http://www.nature.com/articles/nature13005.00162.

[20] P. Jia, L. Wang, A. H. Fanous, X. Chen, K. S. Kendler, Z. Zhao, I. S. Consortium, et al. A bias-reducing pathway enrichment analysis of genome-wide association data confirmed association of the mhc region with schizophrenia. Journal of medical genetics, 49(2):96–103, 2012.

[21] J. M. M. John, T. Cattaert, F. Van Lishout, E. S. Gusareva, and K. Van Steen. Lower-order effects adjustment in quantitative traits model-based multifactor dimensionality reduction. PLoS One, 7(1), 2012.

[22] E. Jorgenson and J. S Witte. A gene-centric approach to genome-wide association studies. Nature reviews. Genetics, 7:885–91, 12 2006. doi: 10.1038/nrg1962.

[23] B. Lehne, C. M. Lewis, and T. Schlitt. From snps to genes: disease association at the gene level. PloS one, 6(6):e20133, 2011.

[24] A. Liberzon, C. Birger, H. Thorvaldsdóttir, M. Ghandi, J. P. Mesirov, and P. Tamayo. The molecular signatures database hallmark gene set collection. Cell Systems, 1(6):417–425, Dec. 2015. doi: 10.1016/j.cels.2015.12.004. URL https://doi.org/10.1016/j.cels.2015.12.004.

[25] Z. Lin, J. P. Hegarty, G. John, A. Berg, Z. Wang, R. Sehgal, D. M. Pastor, Y. Wang, L. R. Harris, L. S. Poritz, et al. Nod2 mutations affect muramyl dipeptide stimulation of human b lymphocytes and interact with other ibd-associated genes. Digestive diseases and sciences, 58(9):2599–2607, 2013.

[26] Z. Lin, Z. Wang, J. P. Hegarty, T. R. Lin, Y. Wang, S. Deiling, R. Wu, N. J. Thomas, and J. Floros. Genetic association and epistatic interaction of the interleukin-10 signaling pathway in pediatric inflammatory bowel disease. World journal of gastroenterology, 23(27):4897, 2017.

[27] J. Z. Liu, S. Van Sommeren, H. Huang, S. C. Ng, R. Alberts, A. Takahashi, S. Ripke, J. C. Lee, L. Jostins, T. Shah, et al. Association analyses identify 38 susceptibility loci for inflammatory bowel disease and highlight shared genetic risk across populations. Nature genetics, 47(9):979, 2015.

[28] L. Ma, A. G. Clark, and A. Keinan. Gene-based testing of interactions in association studies of quantitative traits. PLoS genetics, 9(2), 2013.

[29] T. A. Manolio, F. S. Collins, N. J. Cox, D. B. Goldstein, L. A. Hindorff, D. J. Hunter, M. I. McCarthy, E. M. Ramos, L. R. Cardon, A. Chakravarti, J. H. Cho, A. E. Guttmacher, A. Kong, L. Kruglyak, E. Mardis, C. N. Rotimi, M. Slatkin, D. Valle, A. S. Whittemore, M. Boehnke, A. G. Clark, E. E. Eichler, G. Gibson, J. L. Haines, T. F. C. Mackay, S. A. McCarroll, and P. M. Visscher. Finding the missing heritability of complex diseases. Nature, 461(7265):747–753, Oct. 2009. ISSN 0028-0836, 1476-4687. doi: 10.1038/nature08494.URL http://www.nature.com/articles/nature08494.06874.

[30] D. P. McGovern, J. I. Rotter, L. Mei, T. Haritunians, C. Landers, C. Derkowski, D. Dutridge, M. Dubinsky, A. Ippoliti, E. Vasiliauskas, et al. Genetic epistasis of il23/il17 pathway genes in crohn’s disease dermot. Inflammatory bowel diseases, 15(6):883–889, 2009.

[31] J. H. Moore and S. M. Williams. Traversing the conceptual divide between biological and statistical epistasis: systems biology and a more modern synthesis. BioEssays, 27(6):637–646, June 2005. ISSN 0265-9247, 1521-1878. doi: 10.1002/bies.20236. URL http://doi.wiley.com/10.1002/bies.20236.00327.

[32] C. Niel, C. Sinoquet, C. Dina, and G. Rocheleau. A survey about methods dedicated to epistasis detection. Frontiers in Genetics, 6, Sept. 2015. ISSN 1664-8021. doi: 10.3389/fgene.2015.00285. URL http://journal.frontiersin.org/Article/10.3389/fgene.2015.00285/abstract.00000.

[33] C. Pedros, G. Gaud, I. Bernard, S. Kassem, M. Chabod, D. Lagrange, O. Andréoletti, A. S. Dejean, R. Lesourne, G. J. Fournié, et al. An epistatic interaction between themis1 and vav1 modulates regulatory t cell function and inflammatory bowel disease development. The Journal of Immunology, 195(4):1608–1616, 2015.

[34] S. A. Pendergrass, A. Frase, J. Wallace, D. Wolfe, N. Katiyar, C. Moore, and M. D. Ritchie. Genomic analyses with biofilter 2.0: knowledge driven filtering, annotation, and model development. BioData Mining, 6(1), Dec. 2013. ISSN 1756-0381. doi: 10.1186/1756-0381-6-25. URL http://biodatamining.biomedcentral.com/articles/10.1186/1756-0381-6-25.00042.

[35] A. C. Petrey and A. Carol. The extracellular matrix in ibd: a dynamic mediator of inflammation. Current opinion in gastroenterology, 33(4):234, 2017.

[36] J. Piñero, J. M. Ramírez-Anguita, J. Saüch-Pitarch, F. Ronzano, E. Centeno, F. Sanz, and L. I. Furlong. The DisGeNET knowledge platform for disease genomics: 2019 update. Nucleic Acids Research, Nov. 2019. doi: 10.1093/nar/gkz1021. URL https://doi.org/10.1093/nar/gkz1021.

[37] S. Purcell, B. Neale, K. Todd-Brown, L. Thomas, M. A. Ferreira, D. Bender, J. Maller, P. Sklar, P. I. De Bakker, M. J. Daly, et al. Plink: a tool set for whole-genome association and population-based linkage analyses. The American journal of human genetics, 81(3):559–575, 2007.

[38] K. A. Shaw, D. J. Cutler, D. Okou, A. Dodd, B. J. Aronow, Y. Haberman, C. Stevens, T. D. Walters, A. Griffiths, R. N. Baldassano, et al. Genetic variants and pathways implicated in a pediatric inflammatory bowel disease cohort. Genes & Immunity, 20(2):131, 2019.

[39] X. Sheng and J. Yang. An adaptive truncated product method for combining dependent p-values. Economics letters, 119(2):180–182, 2013.

[40] A. Soroosh, S. Albeiroti, G. A. West, B. Willard, C. Fiocchi, and A. Carol. Crohn’s disease fibroblasts overproduce the novel protein kiaa1199 to create proinflammatory hyaluronan fragments. Cellular and molecular gastroenterology and hepatology, 2(3):358–368, 2016.

[41] A. Subramanian, P. Tamayo, V. K. Mootha, S. Mukherjee, B. L. Ebert, M. A. Gillette, A. Paulovich, S. L. Pomeroy, T. R. Golub, E. S. Lander, and J. P. Mesirov. Gene set enrichment analysis: A knowledge-based approach for interpreting genome-wide expression profiles. Proceedings of the National Academy of Sciences, 102(43):15545–15550, Sept. 2005. doi: 10.1073/pnas.0506580102. URL https://doi.org/10.1073/pnas.0506580102.

[42] D. Szklarczyk, A. L. Gable, D. Lyon, A. Junge, S. Wyder, J. Huerta-Cepas, M. Simonovic, N. T. Doncheva, J. H. Morris, P. Bork, L. J. Jensen, and C. Mering. STRING v11: protein–protein association networks with increased coverage, supporting functional discovery in genome-wide experimental datasets. Nucleic Acids Research, 47(D1):D607–D613, Jan. 2019. ISSN 0305-1048, 1362-4962. doi: 10.1093/nar/gky1131. URL https://academic.oup.com/nar/article/47/D1/D607/5198476.00072.

[43] J. Traherne. Human mhc architecture and evolution: implications for disease association studies. International journal of immunogenetics, 35(3):179–192, 2008.

[44] K. Van Steen and J. Moore. How to increase our belief in discovered statistical interactions via large-scale association studies? Human genetics, 138(4):293–305, 2019.

[45] S. Vermeire, P. Rutgeerts, K. Van Steen, S. Joossens, G. Claessens, M. Pierik, M. Peeters, and R. Vlietinck. Genome wide scan in a flemish inflammatory bowel disease population: support for the ibd4 locus, population heterogeneity, and epistasis. Gut, 53(7):980–986, 2004.

[46] O. A. Vsevolozhskaya, F. Hu, and D. V. Zaykin. Detecting weak signals by combining small p-values in genetic association studies. Frontiers in genetics, 10:1051, 2019.

[47] K. Watanabe, E. Taskesen, A. van Bochoven, and D. Posthuma. Functional mapping and annotation of genetic associations with FUMA. Nature Communications, 8(1), Dec. 2017. ISSN 2041-1723. doi: 10.1038/s41467-017-01261-5. URL http://www.nature.com/articles/s41467-017-01261-5.00139.

[48] W. K. Wu, R. Sun, T. Zuo, Y. Tian, Z. Zeng, J. Ho, J. C. Wu, F. K. Chan, M. T. Chan, J. Yu, J. J. Sung, S. H. Wong, M. H. Wang, and S. C. Ng. A novel susceptibility locus in MST1 and gene-gene interaction network for crohn’s disease in the chinese population. Journal of Cellular and Molecular Medicine, 22(4): 2368–2377, Feb. 2018. doi: 10.1111/jcmm.13530. URL https://doi.org/10.1111/jcmm.13530.

[49] X. Wu, H. Dong, L. Luo, Y. Zhu, G. Peng, J. D. Reveille, and M. Xiong. A novel statistic for genome-wide interaction analysis. PLoS genetics, 6(9):e1001131, 2010.

[50] D. K.-S. Yip, L. L. Chan, I. K. Pang, W. Jiang, N. L. Tang, W. Yu, and K. Y. Yip. A network approach to exploring the functional basis of gene–gene epistatic interactions in disease susceptibility. Bioinformatics, 34(10):1741–1749, 2018.

[51] K. Yu, Q. Li, A. W. Bergen, R. M. Pfeiffer, P. S. Rosenberg, N. Caporaso, P. Kraft, and N. Chatterjee. Pathway analysis by adaptive combination of p-values. Genetic Epidemiology: The Official Publication of the International Genetic Epidemiology Society, 33(8):700–709, 2009.

[52] D. V. Zaykin, L. A. Zhivotovsky, P. H. Westfall, and B. S. Weir. Truncated product method for combining p-values. Genetic Epidemiology: The Official Publication of the International Genetic Epidemiology Society, 22(2):170–185, 2002.

[53] J. Zhang, Z. Wei, C. J. Cardinale, E. S. Gusareva, K. V. Steen, P. Sleiman, and H. Hakonarson. Multiple epistasis interactions within mhc are associated with ulcerative colitis. Frontiers in genetics, 10:257, 2019.

