## supporting information for "Interpretable network-guided epistasis detection"

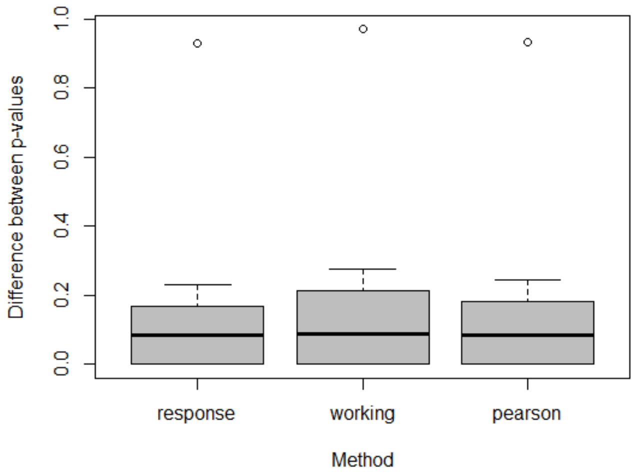


**Supplementary Fig 1:** To choose the best way of computing residuals to obtain the phenotype adjusted for population structure, we randomly extracted five SNPs in the dataset (rs12488468, rs1005678, rs11714286, rs2267844, rs11720964) and compared the associated outputs of epistasis detection. First, we computed the different residuals: we ran a logistic regression model with binary phenotypes as the response variable and 7 PCs as independent variables. We derived three vectors of adjusted phenotypes from the response, working and Pearson residuals. Then, we looked for statistical epistasis: we computed linear models using the different residuals as the response variable and two SNPs and their interactions a independent variables. Finally, we performed logistic regressions with the binary phenotype as the dependent variable, two SNPs and their interaction as explanatory variables, in addition to 7 PCs as covariates. We aimed at identifying the residuals leading to P-values as close as possible to the P-values from the logistic regression. P-values obtained with response residuals as phenotypes are the closest to the ones obtained with the logistic regression and are therefore selected as adjusted phenotypes in our analysis.

**Supplementary Table 1:** The 16 pathways enriched in the *eQTL* analysis. They are obtained from one gene neighborhood (*HYAL1, HYAL3, SPAM1*).

| |
|---|
| GO hyaluronoglycosaminidase activity |
| GO hexosaminidase activity |
| KEGG glycosaminoglycan degradation |
| GO hydrolase activity hydrolyzing o glycosyl compounds |
| GO hydrolase activity acting on glycosyl bonds |
| NABA ecm regulators |
| GO response to UV B |
| GO hyaloronan catabolic process |
| REACTOME hyaluronan metabolism |
| GO hyaluronan metabolic process |
| REACTOME chondoitrin sulfate dermatan sulfate |
| GO aminoglycan catabolic process |
| NABA matrisome associated |
| GO cellular response to UV B |
| REACTOME CS DS degradation |
| REACTOME hyaluronan uptake and degradation |

**Supplementary Table 2:** The 71 pathways enriched in the *Positional + eQTL + Chromatin* analysis. They are obtained from two gene neighborhoods (*HYAL3, HYAL1, HYAL2*, and *PLA2G2E PLA2G5 PLA2G2C*).

| |
|---|
| GO HYALURONONGLUCOSAMINIDASE ACTIVITY |
| GO CELLULAR RESPONSE TO UV B |
| GO HEXOSAMINIDASE ACTIVITY |
| REACTOME HYALURONAN UPTAKE AND DEGRADATION |
| GO RESPONSE TO UV B |
| GO HYALURONAN CATABOLIC PROCESS |
| REACTOME HYALURONAN METABOLISM |
| KEGG ALPHA LINOLENIC ACID METABOLISM |
| KEGG GLYCOSAMINOGLYCAN DEGRADATION |
| GO ARACHIDONIC ACID SECRETION |
| KEGG ETHER LIPID METABOLISM |
| KEGG LINOLEIC ACID METABOLISM |
| GO HYALURONAN METABOLIC PROCESS |
| GO FATTY ACID DERIVATIVE TRANSPORT |
| GO LONG CHAIN FATTY ACID TRANSPORT |
| KEGG ARACHIDONIC ACID METABOLISM |
| GO AMINOGLYCAN CATABOLIC PROCESS |
| GO CELLULAR RESPONSE TO UV |
| KEGG GLYCEROPHOSPHOLIPID METABOLISM |
| KEGG LONG TERM DEPRESSION |

GO HYDROLASE ACTIVITY HYDROLYZING O GLYCOSYL COMPOUNDS

KEGG VEGF SIGNALING PATHWAY

GO FATTY ACID TRANSPORT

KEGG FC EPSILON RI SIGNALING PATHWAY

GO CELLULAR RESPONSE TO LIGHT STIMULUS

KEGG GNRH SIGNALING PATHWAY

GO HYDROLASE ACTIVITY ACTING ON GLYCOSYL BONDS

GO ACID SECRETION

KEGG VASCULAR SMOOTH MUSCLE CONTRACTION

GO ENTRY INTO HOST

chr3p21

GO RESPONSE TO UV

GO MUCOPOLYSACCHARIDE METABOLIC PROCESS

REACTOME GLYCOSAMINOGLYCAN METABOLISM

GO MONOCARBOXYLIC ACID TRANSPORT

NABA ECM REGULATORS

GO CELLULAR RESPONSE TO RADIATION

REACTOME CS DS DEGRADATION

GO AMINOGLYCAN METABOLIC PROCESS

GO INTERACTION WITH HOST

GO CARBOHYDRATE DERIVATIVE CATABOLIC PROCESS

GO CARTILAGE DEVELOPMENT

GO CALCIUM DEPENDENT PHOSPHOLIPASE A2 ACTIVITY

REACTOME ACYL CHAIN REMODELLING OF PI

GO RESPONSE TO INTERLEUKIN 1

chr1p36

GO PHOSPHATIDYLGLYCEROL ACYL CHAIN REMODELING

REACTOME ACYL CHAIN REMODELLING OF PG

GO PHOSPHATIDYLINOSITOL ACYL CHAIN REMODELING

GO PHOSPHOLIPASE A2 ACTIVITY CONSUMING 1 2 DIPALMITOYLPHOSPHATIDYLCHOLINE

GO PHOSPHATIDYLSERINE ACYL CHAIN REMODELING

REACTOME ACYL CHAIN REMODELLING OF PS

GO CONNECTIVE TISSUE DEVELOPMENT

GO PHOSPHATIDYLETHANOLAMINE ACYL CHAIN REMODELING

REACTOME ACYL CHAIN REMODELLING OF PE

KEGG MAPK SIGNALING PATHWAY

GO PHOSPHOLIPASE A2 ACTIVITY

GO RESPONSE TO TUMOR NECROSIS FACTOR

GO RESPONSE TO LIGHT STIMULUS

REACTOME ACYL CHAIN REMODELLING OF PC

GO VIRAL LIFE CYCLE

GO PHOSPHATIDYLCHOLINE ACYL CHAIN REMODELING

GO PHOSPHATIDYLGLYCEROL METABOLIC PROCESS

GO RESPONSE TO VIRUS

GO ORGANIC ACID CATABOLIC PROCESS

GO ORGANIC ACID TRANSPORT

GO CELLULAR RESPONSE TO ABIOTIC STIMULUS

GO RESPONSE TO ANTIBIOTIC

REACTOME METABOLISM OF CARBOHYDRATES

GO PHOSPHATIDYLSERINE METABOLIC PROCESS

REACTOME SYNTHESIS OF PA

---

**Supplementary Table 3:** Tissue-specific *eQTL* and *Chromatin* analyses. For *eQTL*, we took associations from the following tissues: *colon sigmoid, colon transverse, esophagus gastroesophageal junction, esophagus mucosa, esophagus muscularis, pancreas, small intestine terminal ileum, stomach, whole blood, aveALL (average gene expression among ten brain regions), brain anterior cingulate cortex BA24, brain caudate basal ganglia, brain cerebellar hemisphere, brain cerebellum, brain cortex, brain frontal cortex BA9, brain hippocampus, brain hypothalamus, brain nucleus accumbens basal ganglia, brain putamen basal ganglia* and *brain amygdala.* For *Chromatin*, we took contacts measured in: *pancreas, small bowel, orsolateral prefrontal cortex* and *hippocampus.*

| | eQTL | Chromatin |
|---|---|---|
| SNP models (SNPs) | $3 \times 10^5$ (6,769) | 4615 (990) |
| Experimental threshold | $7.3 \times 10^{-7}$ | NA - No epistatic SNP-pair of the permutations is mappable a Biofilter gene-pair |
| Number of significant SNP-pairs (number of SNPs) | 15 (15) | NA |
| Number of significant gene-pairs (number of genes) | 4 (8) | NA |
| Number of significant pathways | 0 | NA |
| Number of components in the gene network | 4 | NA |

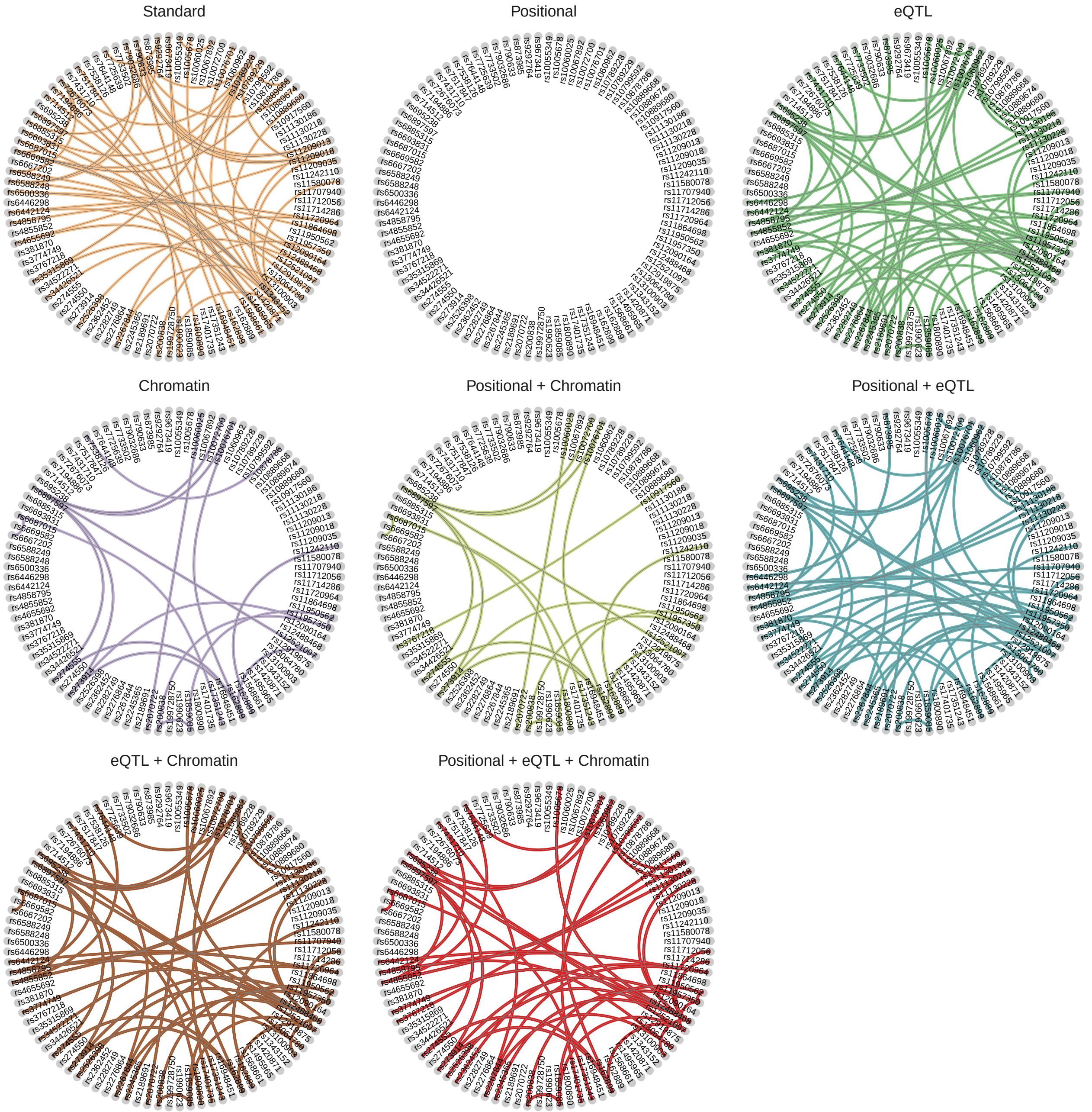


**Supplementary Fig 2:** Epistasis networks built from the significant SNP models of the different analysis.

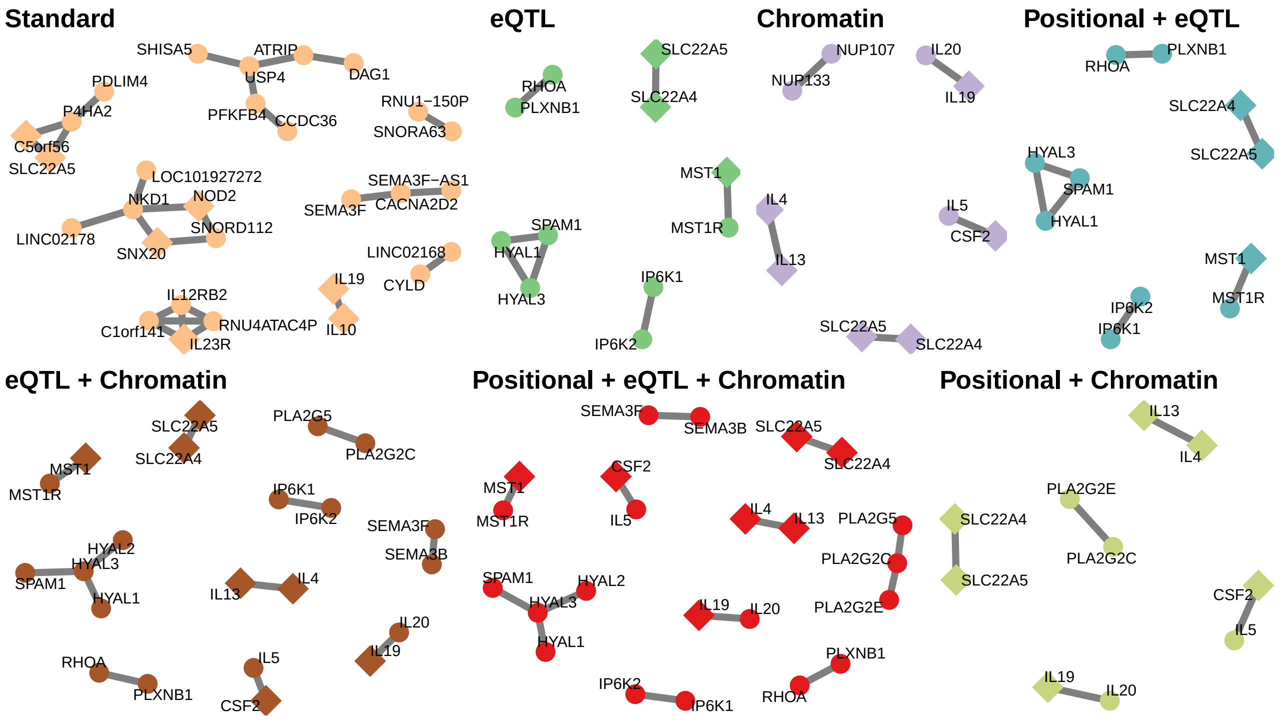


**Supplementary Fig 3:** Alternative visualization of the gene-networks presented in **Figure 2** of the paper. Genes associated to IBD in DisGeNET have a diamond shape.

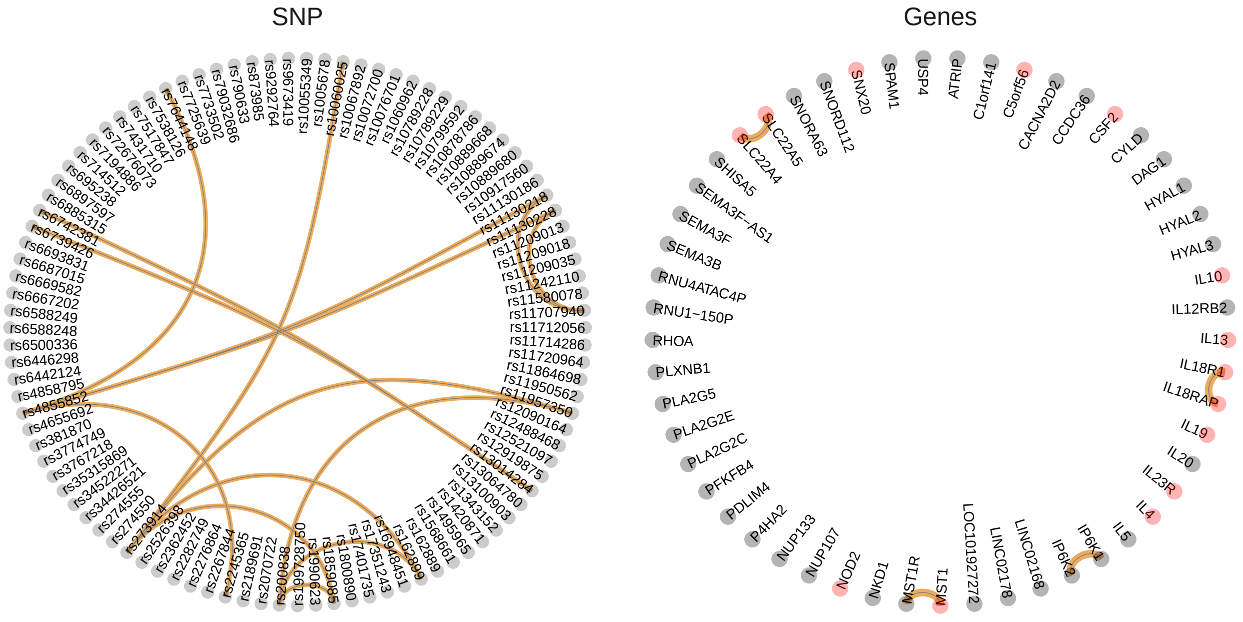


**Supplementary Fig 4:** Epistasis networks built from the significant SNP models and gene models of the eQTL tissue specific analysis. Genes associated to IBD in DisGeNET are colored in pink.
